# Statistical methods for large ensemble of super-resolution stochastic single particle trajectories

**DOI:** 10.1101/227090

**Authors:** N. Hoze, D. Holcman

## Abstract

Following recent progresses in super-resolution microscopy obtained in the last decade, massive amount of redundant single stochastic trajectories are now available for statistical analysis. Flows of trajectories of molecules or proteins are sampling the cell membrane or its interior at a very high time and space resolution. Several statistical analysis were developed to extract information contained in these data, such as the biophysical parameters of the underlying stochastic motion to reveal the cellular organization. These trajectories can further reveal hidden subcellular organization. We present here the statistical analysis of these trajectories based on the classical Langevin equation, which serves as a model of trajectories. Parametric and non-parametric estimators are constructed by discretizing the stochastic equations and they allow recovering tethering forces, diffusion tensor or membrane organization from measured trajectories, that differ from physical ones by a localization noise. Modeling, data analysis and automatic detection algorithms serve extracting novel biophysical features such as potential wells and other sub-structures, such as rings at an unprecedented spatiotemporal resolution. It is also possible to reconstruct the surface membrane of a biological cell from the statistics of projected random trajectories.

## Introduction

Recent advances in super-resolution microscopy [54, 80, 29, 71] associated with single particle tracking (SPT) [73, 75] have allowed routine collections of tens of thousands of short and long trajectories. These large datasets offer new challenges in analysis, interpretation, search and reconstruction of hidden features, especially in the context of cell biology probed at the molecular resolution. These trajectories explore part of a cell, either in dimension 2 [26, 62] or 3 at the shortest possible resolution, and their paths are critically influenced by local organization, as emphasized in neuronal cells [76, 68]. Thus, a large research effort in that direction has been based on the hypothesis that it would be possible to reconstruct some of the biological nano- to micro-structures from these trajectories. It should also be possible to unravel their underlying physical processes.

In the past recent years, these challenges of cell membrane reconstruction, structural identification and membrane organization have been addressed following the development of novel statistical methods based on stochastic models. These models are the Langevin equation, its Smoluchowski limit and associated models that account for additional localization point identification noise. We present here the development of these statistical approaches that are based on stochastic models, the deconvolution procedure and the numerical simulations used to extract biophysical parameters from single particle trajectories data. Several empirical estimators have been proposed to recover the local diffusion coefficient, vector field and even organized patterns in the drift, such as potential wells.

For studying long trajectories, the most common procedure consists in estimating the mean-square displacement (MSD) of the particle [19]. The MSD is proportional to time and allows in theory estimating the diffusion coefficient. It has been widely used in early applications of long but not necessarily redundant single particle trajectories in a biological context [64, 74]. However, the MSD applied to long trajectories suffers from several issues. First, it is not precise in part because the measured points are correlated [87]. Second, it cannot be used to compute any physical diffusion coefficient when trajectories consists of switching episodes for example alternating between free and confined diffusion, where other algorithm procedures have been developed [27]. At low spatiotemporal resolution of the observed trajectories, the MSD behaves sublinearly with time, a process known as anomalous diffusion [84, 56, 58], which is due in part to the averaging of the different phases of the particle motion. To overcome these issues and distinguish between different types of motion, a stochastic model of the acquired data is needed.

This manuscript is organized as follow: we first describe single stochastic particle trajectories recorded in the context of cell biology in two dimensional membrane and in the cell cytoplasm. In the second part, we introduce Langevin stochastic processes used as a model to interpret physical trajectories. These equations are a key step for the construction of parametric and non-parametric empirical estimators to extract forces, diffusion tensor or membrane organization from measured trajectories. The third part explains how suborganization such as potential wells are reconstructed from trajectories. The fourth section shows how the error in the position localization affects the first and second moments computed from SPT trajectoriess. In section five, we present a different type of SPT trajectories analysis used to recover tethering forces applied on particle belonging to a polymer chain (model of a DNA chromatin locus). In the last section, we show how to reconstruct a surface from the projections of many SPT trajectories. This presentation is intended for statisticians, applied mathematicians, theoretical physicists and more general anybody interested in novel methods based on stochastic processes, recently used to extract information from super-resolution data below the diffraction limit.

## 1 Presentation of single particle tracking trajectory data in cellular biology

Single particle trajectories can be short and long often acquired from digital images [32, 54]. Before explaining how they are used to investigate the local structure of the environment they sample, we shall present the trajectories in the context of statistical analysis of these data. Indeed, the goal of building a statistical ensemble from these data is to observe local physical properties of the particles, such as velocity, diffusion, confinement or attracting forces reflecting the interactions of the particles with their environment. It might even now be possible to use modeling to construct from diffusion coefficient (or tensor) the confinement or local density of obstacles reflecting the presence of biological objects of different sizes.

In the context of cell biology single particle trajectories are acquired on tagged molecules or proteins that are not immobile but are usually moving inside the cell either on the two dimensional surface (membrane) or inside the three dimensional cytosol. In all cases, they can be sampled at a nanometer scale of molecular motion. It is possible to observe proteins of interest because they have been genetically or directly associated with a fluorescent molecule that can be tracked under the microscope. The luminescent property of the dye defines their stability and thus the time they will remain bright before they bleach. Collecting these data requires to define a spatial precision, which is usually limited by the diffraction limit [69] and the sampling time acquisition.

Biophysical and molecular processes live at the nanometer scale, that was unaccessible due to the limitation of the spatiotemporal resolution, and the image of a infinitely small particle could not be resolved below hundreds of nanometers (Abbe’s diffraction limit [52]). Detecting objects under the diffraction limit is now routinely achieved using different microscopy techniques such as PALM [7], STED [31] or STORM [69]: these methods rely on stochastic activation of isolated particles (molecules of interest) for a short time. Not all particles emit light at the same time, thus it becomes possible to distinguish their position. For all microscopy approaches, the localization method consists in fitting a Gaussian to the illumination profile of any localized point. The standard deviation of the fitted Gaussian, which determines the precision of localization, is reduced when the emitting particle is of small size, the number of photons received during the acquisition is large and the particle motion is limited.

SPT allowed observing moving particles at the micrometer scale, to investigate cell membrane organization [73, 49, 64, 18, 74, 32, 25], but also the cell nucleus dynamics and mRNA production [22, 81]. Due to the constant improvement of the instrumentation, the spatial resolution is continuously decreasing, reaching today values of approximately 20 nm, while the acquisition time step is usually in the range of 10 to 50 ms to capture the shortest events in live tissues. Once points are acquired, the second step is to reconstruct trajectories. This is done by tracking algorithms that allow connecting the dots of acquired points [14, 75].

We now present trajectories of different moving molecules obtained from sptPALM and other techniques (Fig. 1): First, proteins trafficking on the surface of neuronal cells (transmembrane proteins) [26, 62, 53] can now be recorded at a spatial resolution of approximately 30 nm and a time resolution of 20 ms in sub-surface compartments, such as the pre-synaptic membrane [76, 68]. More recently, sptPALM was used to detect the dynamical organization of membrane proteins in plant cells [36, 47], or events of DNA binding by transcription factors in mammalian nucleus [24] and by CRISPR-Cas9 [46]. The physical interpretation of the trajectories depends on the quality of the upstream data acquisition. Although superresolution image acquisition and particle tracking are crucial to guarantee high quality data, we leave these topics outside as there have been already discussed in [15, 52].

**Figure 1:**
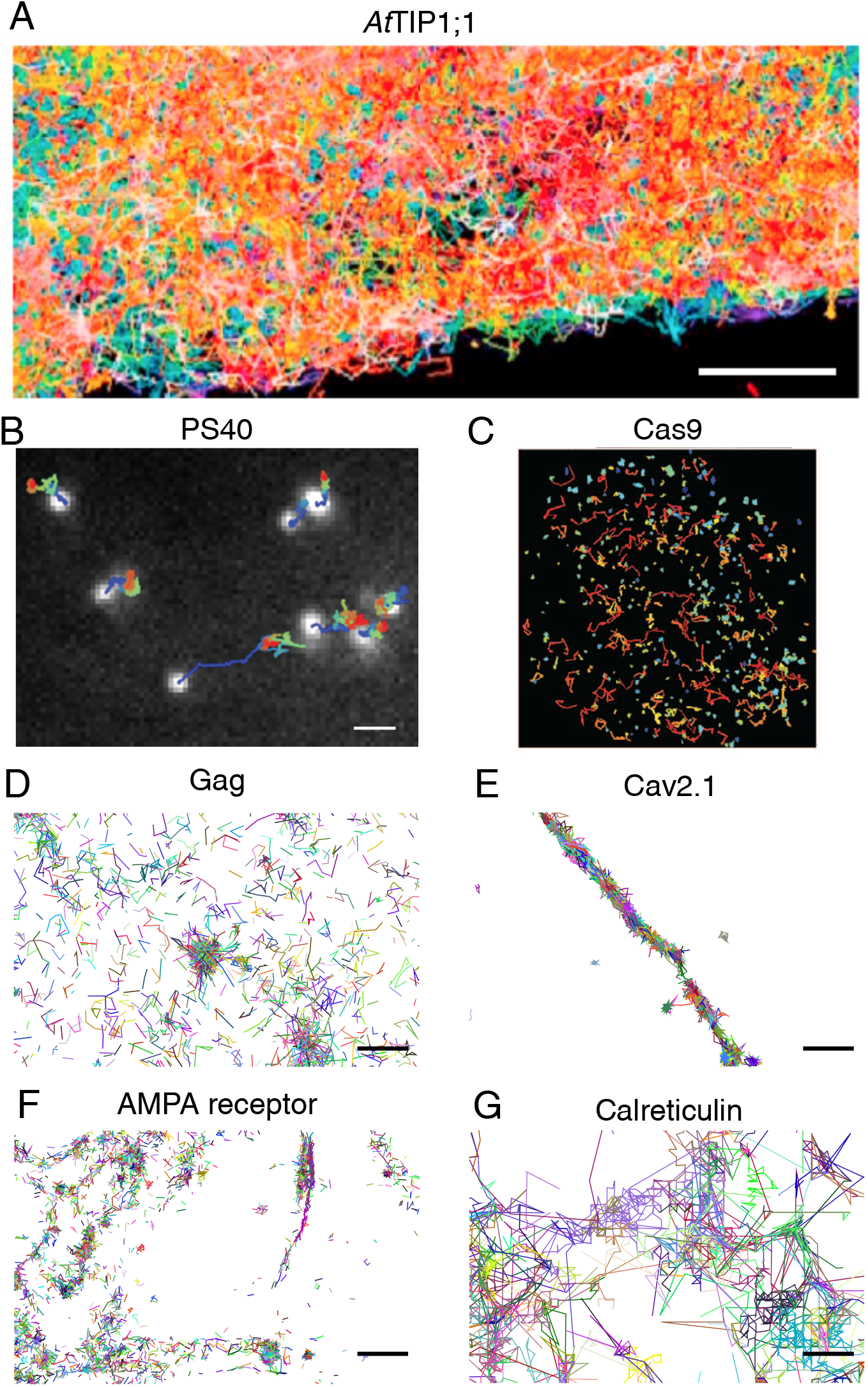
Examples of superresolution trajectories. (A) Superresolution trajectories of the tonoplast marker *At*TIP1;1 in root epidermal cells (adapted from [36]). (B) Interaction of the nanoparticle PS40 with clathrin-coated pits on the plasma membrane of COS-7 cells (adapted from [51]). (C) Dynamics of Cas9 molecules transfected with short interspersed nuclear elements (SINE) of the B2 type in 3T3 cells (adapted from [46]). (D) HIV structural protein Gag at the surface of cellular plasma membrane. (E) Calcium channel Cav2.1 in the presynaptic density. (F) AMPA receptors in hippocampal neurons. (G) Calcium storage protein Calreticulin in the endoplasmic reticulum. Scale bars: A: 5 *μ*m, B, D–G: 1 *μ*m.

## 2 Stochastic equation as a model of motion

### 2.1 Langevin and Smoluchowski equations

Langevin’s equation [19, 8, 50] describe a stochastic particle driven by a random force and a field of force (e.g., electrostatic, mechanical, etc…). With an external force *F*(*x,t*) [13], it is written as

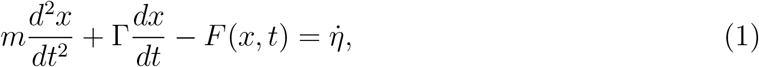

where *m* is the mass of the particle and Γ = 6*πaρ* is the friction coefficient of a diffusing particle, *ρ* the viscosity and *η* the *δ*-correlated Gaussian white noise [19, 8, 50]. The force can derive from a potential, *F*(*x,t*) = –*U′*(*x*) and in that case, with notation *ε* = *k_B_T*, which represents the energy (*k_B_* is the Boltzmann’s constant and *T* the temperature), the Langevin’s equation is transformed into the two dimensional stochastic system

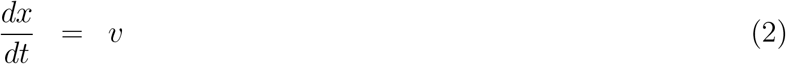

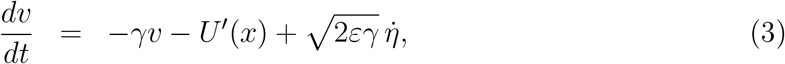

where *γ* = Γ/*m* is the dynamical friction coefficient per unit mass.

Langevin’s equation is used to describe trajectories where inertia or acceleration matters. For example at very short timescales, when a molecule unbinds from a binding site, it escapes the potential well and the inertia term allows the particles to move away from the attractor and thus prevents immediate rebinding that could plague numerical simulations. In the large friction limit *γ* → ∞ the trajectories *x*(*t*) of the Langevin equation 1 converges in probability to these of the Smoluchowski equation [77, 5]

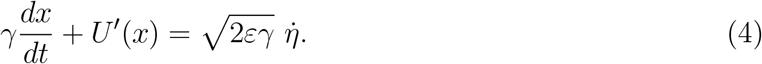

Equation 4 is written equivalently as

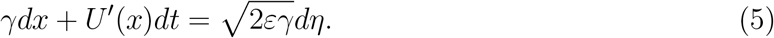

### 2.2 Physical model equations

For a timescale much longer than elementary molecular events, the position of tracked particles is described by the overdamped limit of the Saffman-Delbrück-Langevin model [70, 50]. The model is described in the previous subsection and assumes that the diffusion of a protein or a particle embedded in a membrane surface is generated by a diffusion coefficient *D* and a field of force *F*(*X,t*), according to the overdamped Langevin equation

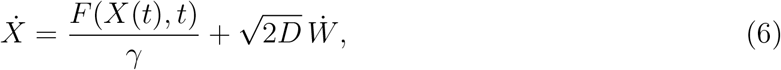

where *W* is a Gaussian white noise and *γ* is the dynamical viscosity [77]. The source of the driving noise is the thermal agitation of the ambient lipid and membrane molecules. However, due to the acquisition timescale of empirical recorded trajectories, which is too low to follow the thermal fluctuations, rapid events are not resolved in the data, and at this coarser spatiotemporal scale, the motion is described by an effective stochastic equation [37, 40]

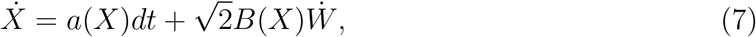

where *a*(*X*) is the drift field and *B*(*X*) the diffusion matrix (see Fig. 2). The effective diffusion tensor is given by 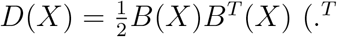 denotes the transposition) [78, 77]. The observed effective diffusion tensor is not necessarily isotropic and can be state-dependent, whereas the friction coefficient *γ* in 6 remains constant and the microscopic diffusion coefficient (or tensor) may remain isotropic.

**Figure 2:**
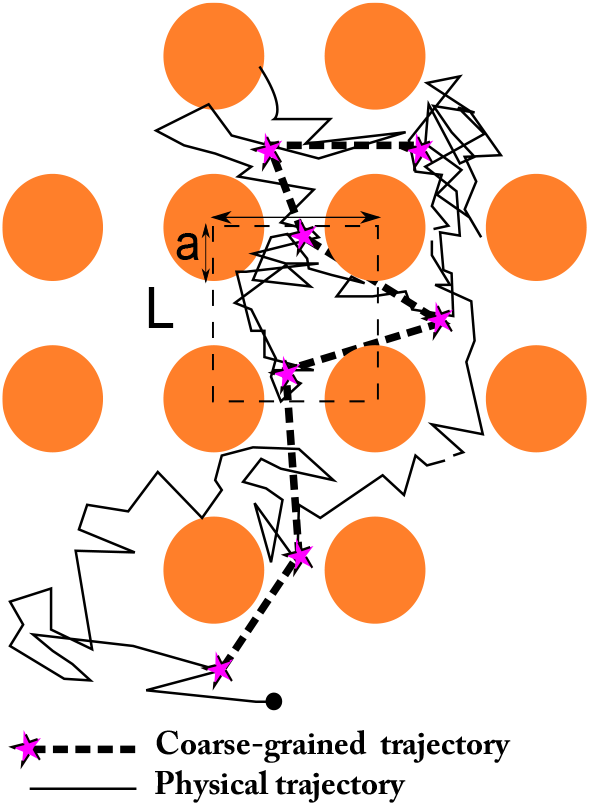
Scheme of a coarse-grained trajectories from an effective stochastic equation. A physical trajectory is generated by the overdamped Langevin equation (Eq. 6) while the coarse-grained trajectory is sampled at fixed time interval Δ*t* (indicated by purple stars). The presented obstacles are not directly visible.

### 2.3 Empirical estimation of the drift and diffusion tensor of a general stochastic process

In this section, we present the construction of empirical estimators that serve to recover physical properties from parametric [37, 41, 87] and non-parametric statistics [4]. Retrieving statistical parameters of a diffusion process from one-dimensional time series statistics have been studied using Bayesian inference in [30, 65]. The approach we present here is based on the direct asymptotic of stochastic equations to construct direct empirical estimators for recovering drift and diffusion tensor. There are derived from discretizing the stochastic equation 7 [44, 78, 79, 33, 34].

We note that the models and the analysis presented above assume that processes are stationary, so that the statistical properties of trajectories do not change over time. In practice, this assumption is satisfied when trajectories are acquired for less than a minute, where only few slow changes may occur on the surface of a neuron for example. Time-lapse analysis [37] with a delay of 15 minutes between acquisitions has indeed revealed slow changes over time of biological significance.

The coarse-grained model 7 is recovered from the conditional moments of the trajectory increments Δ*X* = *X*(*t* + Δ*t*) – *X*(*t*) (see [77, 23, 82]),

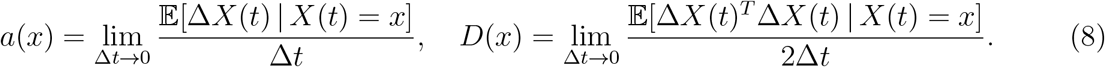

Here the notation 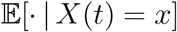 means averaging over all trajectories that are at point *x* at time *t*. Indeed, the coefficients of eq. 6 can be statistically estimated at each point of the membrane (or rather in the focal plane) from an infinitely large sample of its trajectories in the neighborhood of the point *X* at time *t*. In practice, the expectations in 8 are estimated by finite sample averages and Δ*t* is the time-resolution of the recording of the trajectories, as described in [37]. Formulas 8 are approximated in [37] at the time step Δ*t* = 50 ms, where 200 points falling in any bin is usually enough for the estimation.

To estimate the local drift and diffusion coefficients, the trajectories are first grouped within a small neighbourhood. The field of observation is partitioned into square bins *S*(*x_k_, r*) of side *r* and centre *x_k_* (Fig. 3A), and the local drift and diffusion are estimated for each of the square. Considering a sample of *N_t_* trajectories {*x^i^*(*t*_1_),…, *x^i^*(*t_N_s__*)}, where *t_j_* are the sampling times, the discretization of equation 8 for the drift *a*(*x_k_*) = (*a_x_*(*x_k_*), *a_y_*(*x_k_*)) at position *x_k_* is

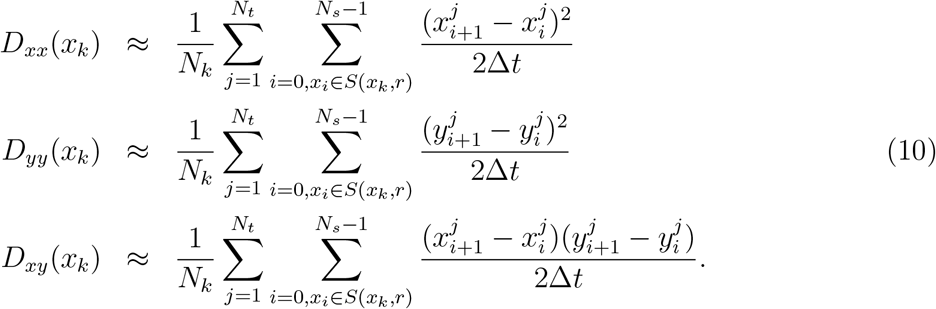

where *N_k_* is the number of points of trajectory that fall in the square *S*(*x_k_, r*). Similarly, the components of the effective diffusion tensor *D*(*x_k_*) are approximated by the empirical sums

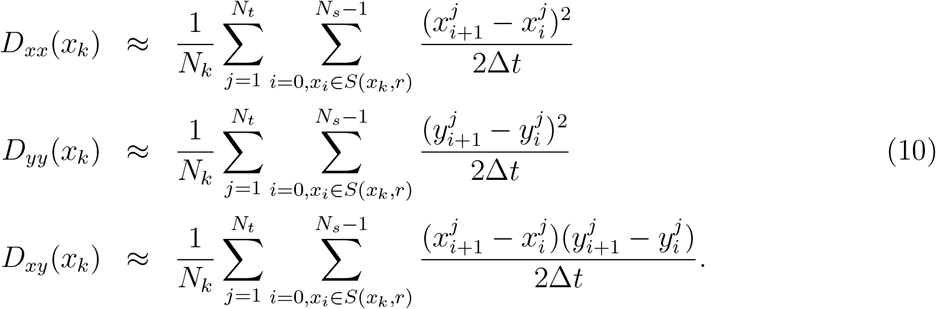

**Figure 3:**
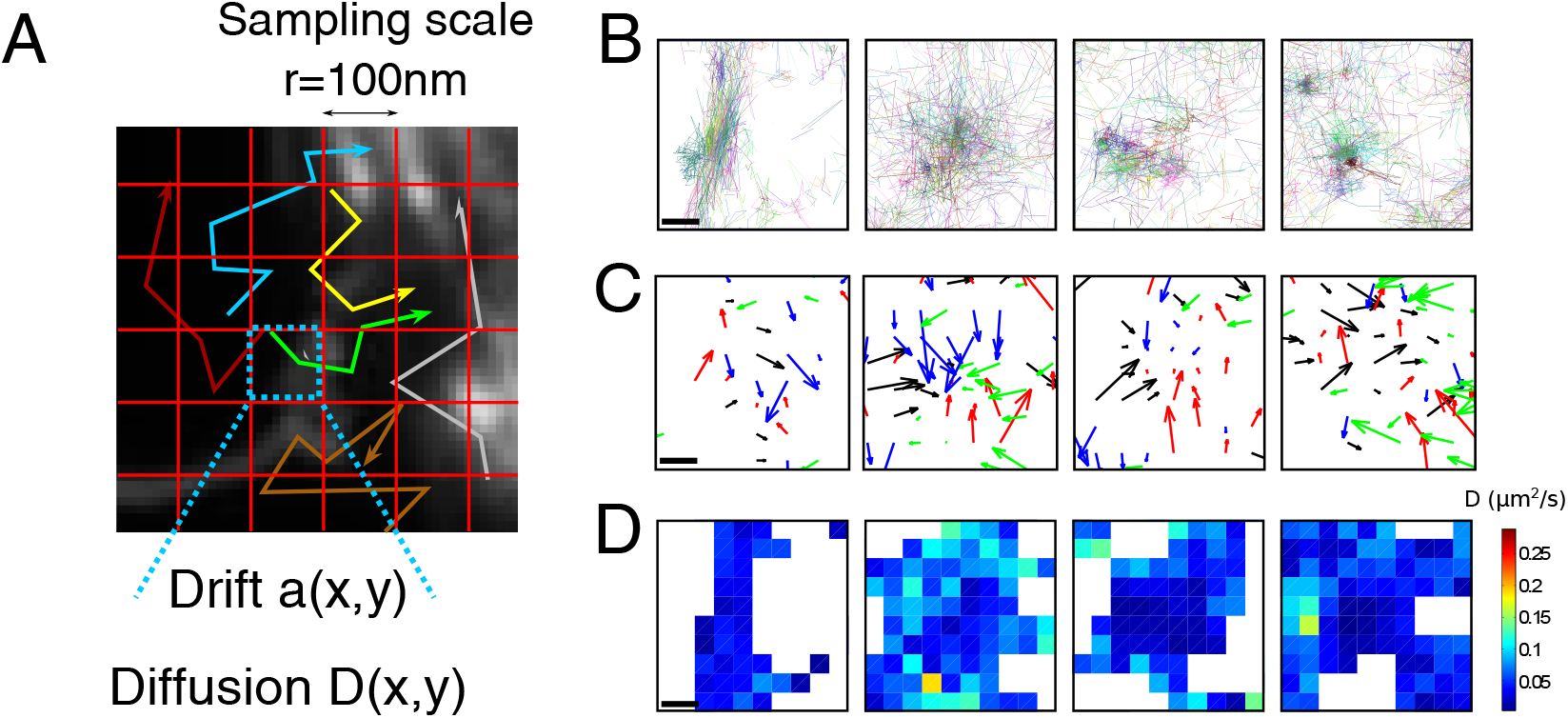
(A) Discretization of the image into pixels of size *r*, that is used preliminarily to the estimation of the local drift and diffusion (eqs. 9–10). (B-D) Application of the estimators to VSVG superresolution trajectories. (B) Four samples of sptPALM trajectories (*n*=30,000) of VSVG proteins. (C) Drift field estimation in the four squares. The arrows are colored depending on the drift orientation. Square side is *r* = 100nm. (D) Map of the diffusion coefficient. Scale bars: 200 nm.

The moment estimation 9 and 10 requires a large number of trajectories passing through each point of the surface, which fits precisely to the massive data generated by the sptPALM technique on biological samples. Indeed, the exact inversion formula eq8 demands in theory an infinite number of trajectories passing through any point *x* of interest. In practice, the recovery of the drift and diffusion tensor is obtained after a region is subdivided by a square grid of radius r (of the order of 50 to 100 nm). Estimations of the error term in the drift compared to the diffusion coefficient suggest a minimum of 200 points per bin (Supp. Information in [37]).

We illustrate this method on trajectories of vesicular stomatitis virus G protein (VSVG) (Fig. 3B). The trajectories are separated on squares of size *r* = 100 nm. The empirical estimator 9 is applied to the trajectories to compute the drift from the trajectories (Fig. 3C). The average diffusion coefficient 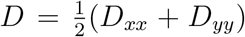 is estimated using eqs. 10 (Fig. 3D).

## 3 Interpretation of two-dimensional regions of high trajectories density as potential wells

The drift field *a*(***x***) in equation 7 represents a force that acts on the diffusing particle, regardless of the existence or not of a potential well [38]. In the case where ***D***(***x***) is locally constant and the coarse-grained drift field ***b***(***x***) is a gradient of a potential

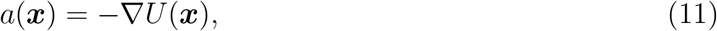

then the density of particles represents locally the Boltzmann density *e*^−*U*(***x***)/*D*^ [34]. The force field can form potential wells, generically approximated locally as a paraboloid with an elliptic base. It remains a difficult question to extract automatically the axis, the center and the boundary of the elliptic base of the well. Once they are known, within the analytical representation 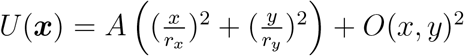, the constants *A, r_x_, r_y_* are three parameters to be determined.

We present methods to detect empirical potential wells reconstructed from data. In particular, several parameters influence the detections such as the pixel resolution and the minimum number of sampled points of trajectory in each pixel. Another interesting feature is the time evolution of these structures.

### 3.1 Vector fields, attractors and characteristic index

A determinist vector field is most of the time a local non-zero flow. However, there are rare occasions where it vanishes at isolated points, and each region near these points is of particular interest. The situation is however more complicated when the vector field has random components, especially near critical points, because the noise dominates. Yet, to determine whether a vector field of the extracted drift component contains structures such as basin of attractions of point attractors (which is probably the simplest detectable structure of a generic dynamical system), a robust characterization of the field is needed to differentiate random distributions from those that may contain a physical significance. Due to the time and space resolution limitation, in the context of SPT, small attractors, associated with short-range interactions less than few tens of nano-meters, below the diffraction limit, cannot be resolved.

To characterise a two-dimensional attractor, generated by a gradient field, the first step it to recover the elliptic paraboloid, which is second order Taylor approximation of the field at the non degenerated critical point. The elliptic domain is the basin of attraction, while its depth is a measure of the force field strength. For a circular potential well of depth *A* and radius *r*, centered at a point (*x*_0_, *y*_0_), the field is given by

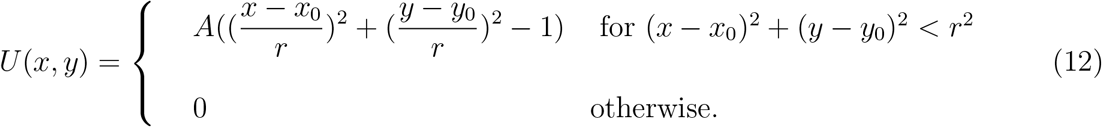

The associated field of force with a critical point at the center is an ubiquitous pattern of interaction (see Fig. 4A-B). The nature of potential wells in empirical data is still unclear [33] as they can either be due to a direct interaction of a receptor, proteins or a molecule with a partner such as a scaffolding protein or with a coherent ensemble of organized molecules.

**Figure 4:**
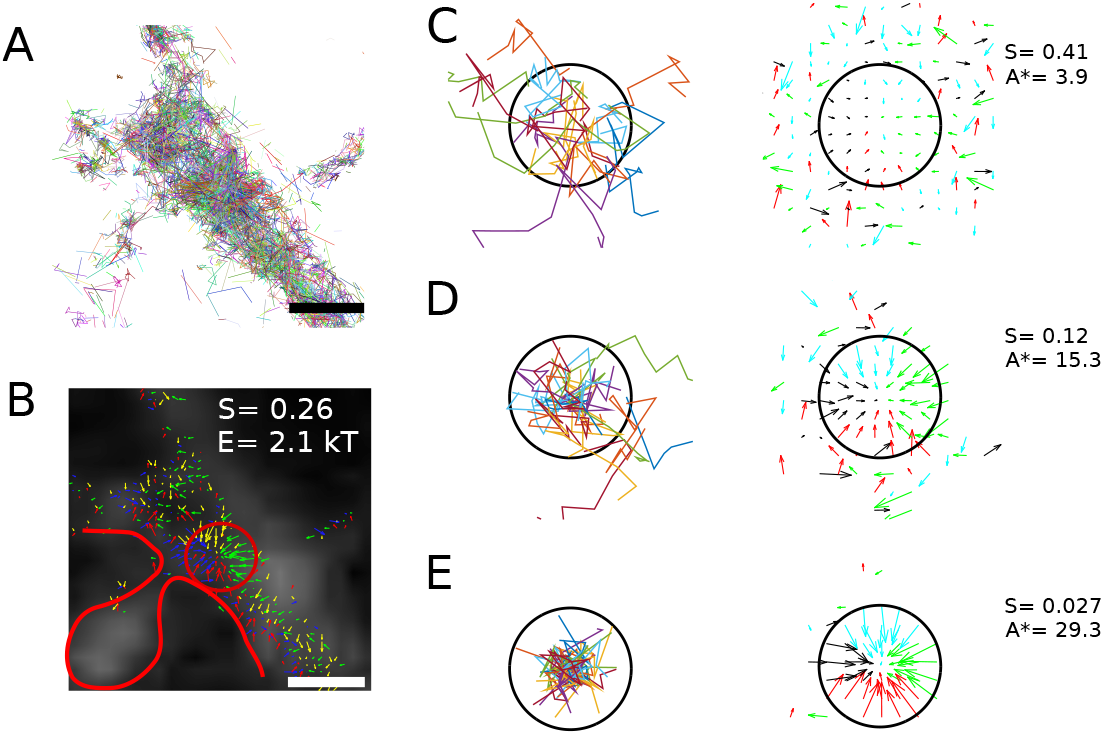
Potential well strength estimation. (A) Trajectories of AMPA receptors in a high density region, at the base of a dendritic spine: characterized by a potential well (B), as indicated by converging a vector field (adapted from [34]). Scale bar: 1*μ*m. (C,D,E) Analysis and estimation of potential well strength on simulated trajectories. The process stochastic is 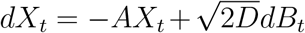, inside a disk of radius 1 (black), and pure diffusion 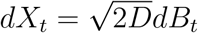 outside. Twenty trajectories, each containing ten points are shown (left) and the drift map is estimated over 400 trajectories, using formula 9. Parameters are *D* = 1*μ*m^2^/*s*, Δ*t* = 0.05*s*, and (C): *A* = 2, (D): *A* = 10, (E): *A* = 20. The index *S* and the estimated *A* are indicated for each set of trajectories. Here *R* =1 (no units), bins are 0.3, so there are about 36 bins per potential well.

To recover the strength *A* of a potential well from a spatial drift distribution (eq. 9), a minimization procedure was introduced [37] based on evaluating the error between the field of force and the closed gradient:

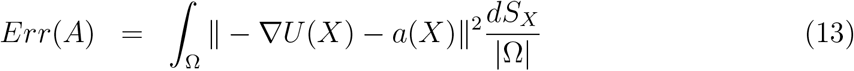

where Ω is the basin of attraction. In practice, using a sampling set of points *X*_1_,‥, *X_N_*, after discretizing the error 13, we obtain

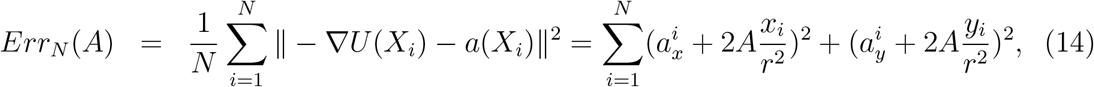

where 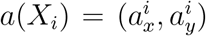 is the empirical vector field on the square centered on *X_i_*, and we set (*x*_0_, *y*_0_) = (0, 0) for the sake of simplicity. The harmonic potential that minimizes the distance to the field *a* is attained for the value *A*^*^ given by

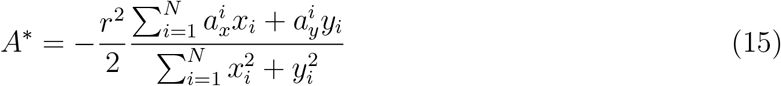

and the minimal error is

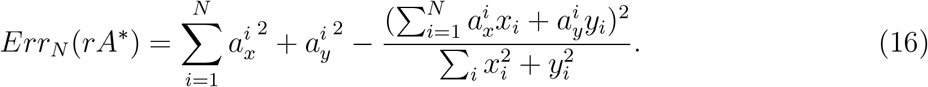

The distance between a vector field and one derived from a harmonic potential well in its basin of attraction is characterized by the index

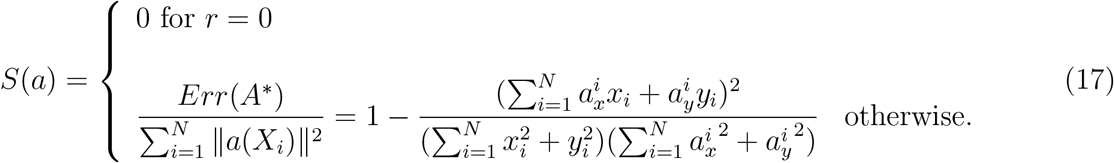

The index *S*(*a*) varies in the interval [0,1]. For a pure harmonic potential *S*(*a*) = 0, while a drift made of independent Brownian direction (no potential) is characterized by lim_*N*→∞_ *S_N_* = 1. This minimization procedure is illustrated in Fig. 4C-E, where the drift is computed from simulated trajectories for potential wells with various strengths. The estimation of the well strength depends on the number of trajectories. The index can be used for elliptic and not only circular wells.

To conclude, empirical estimators are constructed from the first and second moment formula 8. These estimators have been applied to extract features from SPT trajectories [34, 40, 37]. Converging arrows have been interpreted as potential wells, but their biophysical nature remains unclear [33]. They could be due to membrane curvature, direct interactions with organized scaffolding proteins, local receptor interactions with converging cytoskeleton or a combination of all of them. The energies of the wells for membrane receptors or channels are distributed between 1.5 to 8 *k_B_T*, although it is slightly reduced for voltage-dependent calcium channels than for AMPA receptors [63, 34]. In the next section, we will show how the localization noise impacts the recovery of the first moments and the extraction of the potential wells. Finally, potentials wells can be organized in ring domains as reported in [38].

## 4 Model of a stochastic process perturbed by error localisation noise and parameter estimations

The statistical methods now assume that SPTs are already reconstructed from an ensemble of points, estimated from tracking algorithms [14]. There are several sources of the noise: the acquired points are corrupted by errors approximated as Gaussian. This simple model accounts for localization errors when the emitters (dye particles) have a constant intensity during the trajectory acquisition and in addition, the noise source is identical for all trajectories. This noise is associated to localization errors introduced by assigning a random position of the detected particle within the Gaussian localization point spread function. This reconstruction procedure is now a classical for tracking and implemented in freely available softwares [14].

However, we do consider here errors originating from an upstream tracking algorithm that mistakenly combines spots from two different molecules. This error can potentially bias the reconstruction of trajectories, but as different noisy time series are combined when trajectories visit the same spatial landmarks in the cell, this error may actually be smoothed out by the averaging procedure based on formula 8.

Finally, we emphasize that statistical issues inherent to approximation for transition densities and time series estimation have been studied in the context of classical statistics, but we emphasize here complications due to small time approximation in the moment estimators [65, 1, 30, 48]. We shall present in this section some singular behaviors when the Gaussian measurement noise corrupts the signal. Thus the approach we present here takes the point of view of analytical approximation and statistical physics.

To reconstruct a diffusion process from trajectories that contains additional errors introduced by tracking algorithms or uncertainty in the detection process [85], a model of these errors is necessary. Indeed, the exact localization of the trajectory points is affected, thus modifying the estimation of the biophysical parameters such as the diffusion coefficient. The different noise terms not only bias the parameter estimates, they also induce a time correlation between the observed points of the trajectories. In this section, we present the possible noise sources and the models developed to take these fluctuation noises into account [6, 87].

A model of the localization noise introduced in measuring the position of a particle is obtained by fitting a Gaussian on the diffraction spot. The localization uncertainty accounts for the width of the Gaussian standard deviation *σ*, the signal intensity, the intrinsic fluctuation of individual photons and the background noise [85]. In a first approximation, the observed position *y_n_* at time *t_n_* = *n*Δ*t* is then obtained from the true position *x_n_* = *x*(*t_n_*) by adding a Gaussian variable *ξ_n_*

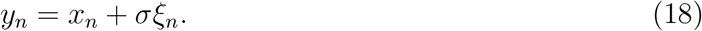

Another source of errors originates from the finite camera exposure, which is due to the fact that collected photons are coming from a moving stochastic particle. The choice of the detected point results from the integration of the particle positions between two acquisition time points. This additional noise, called the motion blur [6], dynamic errors [72] or dynamic localization uncertainty [59] depends on the number of acquired photons and on the displacement of the particle between two time intervals [59]. The motion blur can influence the point spread function resulting in breaking symmetry and non Gaussian noise description. In this section, we consider only a Gaussian localization noise, which is the same for all particles at any acquisition time step. This assumption is discussed in [12], where a time-dependent localization noise is chosen.

The camera shutter state function *ς*(*t*) characterizes the opening of the camera during a time interval Δ*t*. The camera shutter state affects the observed position *x_true_* of the particle at time *t_n_* = *n*Δ*t* and leads to the choice of the position

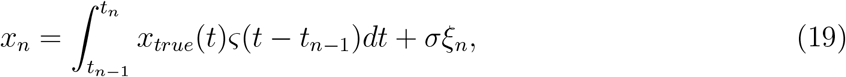

where

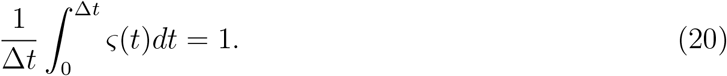

The motion blur coefficient is then given by

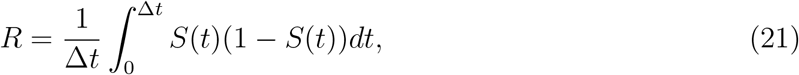

where 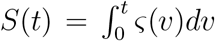. Recovering *x_true_* from the sequence *x_n_* and estimating the physical diffusion coefficient involves computing various statistical moments such as the mean square displacement. However, this estimation needs to account for the various sources of noise [6, 87, 88]. Indeed, the localization error and finite camera exposure affect the MSD [72, 61, 28], and the motion blur coefficient appears as a correction term in the diffusion coefficient estimation. For example, the finite exposure time critically affects the estimate of the diffusion coefficient for particles undergoing hop diffusion or confined motion [67] and the measured coefficient *D* decreases with the exposure time. The motion blur has higher impact on fast moving particles, due to integration over longer areas. We present now some empirical estimators.

### 4.1 Local estimators applied to noisy trajectories

Estimating drift and diffusion with a localization noise from the empirical estimators 9 and 10 supposes that the physical stochastic process [41] is

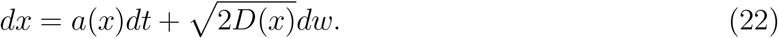

This section relies on the short time approximation

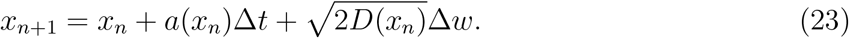

Euler’s numerical scheme of stochastic equation 23 and approximations of the transition probability density function are commonly used in the context of statistical inference of a diffusion process, time series analysis and financial econometrics literature [65, 1, 30]. Here, we discuss only an asymptotic approach to estimate the first and second moments of a stochastic process and the two-dimensional vector field underlying the drift term. The asymptotic approach provide an estimation of the estimators at order *o*(Δ*t*).

**Figure 5:**
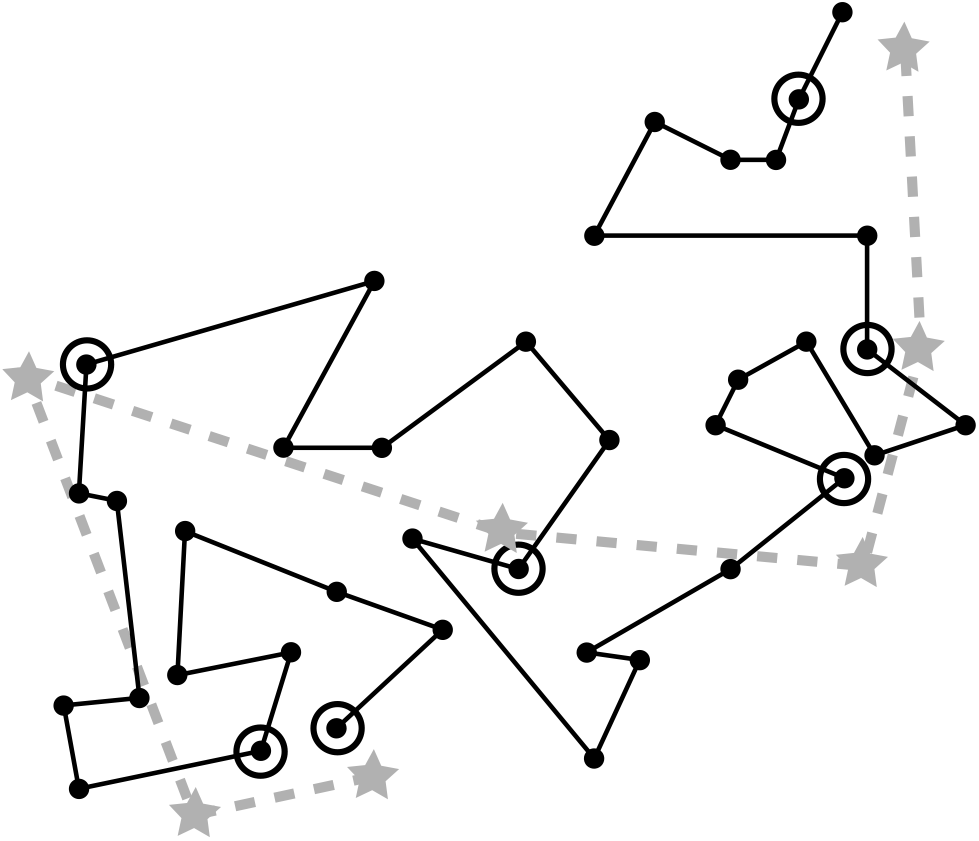
Observed trajectory (grey dashed line) sampled from a physical process (black line) and perturbed by a localization noise. At constant time interval Δ*t*, the physical trajectory is sub-sampled (black circles). Localization noise perturbs the localization and the observed points (grey stars) are positioned in a neighborhood of physical ones.

We start when the observed points are at positions *x_n_* for the physical process, generated by relation 18. The deconvolution of the trajectories consists in removing the Gaussian localization noise from the measured trajectories acquired at a time resolution Δ*t*. An observed trajectory is a sequence of points *y_n_* sampled from a physical process *x_n_* and corrupted with a Brownian noise *σ_n_*, where *σ* > 0 and *ξ_n_* are i.i.d Gaussian variable of variance 1 (relation 18). We neglect the motion blur coefficient (*R* = 0). To compute the first moment, each component in relation 23 are expanded each as follow:

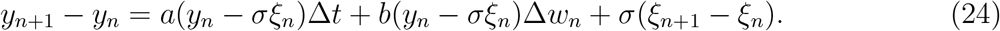

Using a Taylor expansion

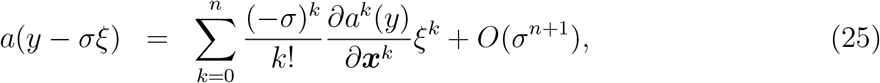

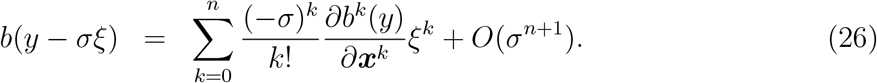

at second order (in dimension one),

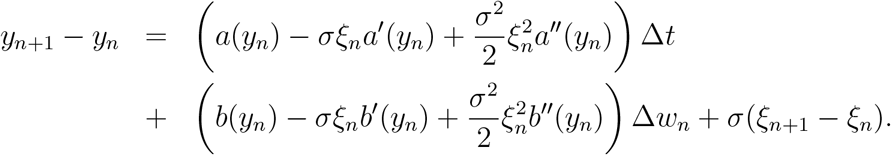

the expectation is computed with 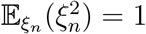

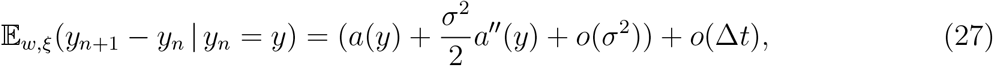

We conclude that at order two, a correction has to be added to the drift, but when *σ* is small this contribution is negligible. In particular, this result shows that at the first order the additive noise does not influence the recovery of the vector field and local potential wells. The recovered energy is thus not affected by additive noise. The diffusion coefficient is computed from the second moment for Δ*t* ≪ 1,

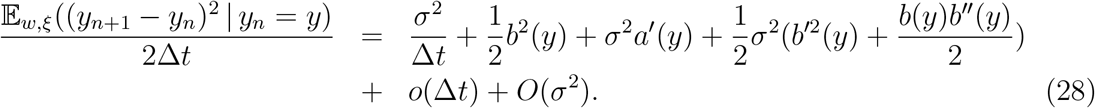

The correlation term with the drift comes from the product –*σξ_n_a′*(*y_n_*)(*σ*(*ξ*_*n*+1_ – *ξ_n_*)). Due to the independent property of the random variables, the product 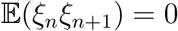, but 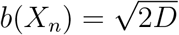. We are thus left with new term *σ*^2^*a′*(*y*) in the effective diffusion coefficient, in agreement with the numerical simulations.

Another method to compute the statistical moment uses the transition probability *p_n_*(*y*|*x*) = *Pr*{*y*_*n*+1_ = *y*|*y_n_* = *x*} of the observed point at position *y* and at time (*n* + 1)Δ*t* given that at time *n*Δ*t* it is observed at position *x*, is computed from the pdf of the physical process by a convolution with the Gaussian distribution of the localization noise. In one dimension, when the diffusion tensor 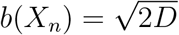 is uniform in space

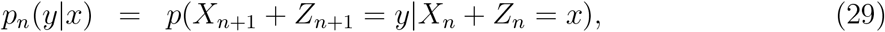

where the two processes *X_n_* and *Z_n_* are independent. In 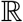, we have

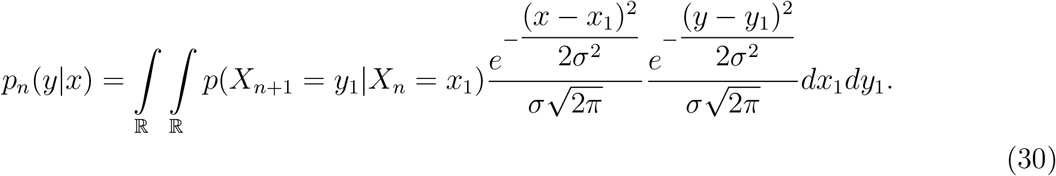

For Δ*t* ≪ 1, we use the approximation of the 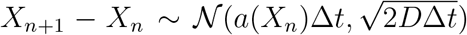. The transition probability is thus approximated by 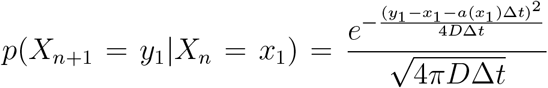, which gives that 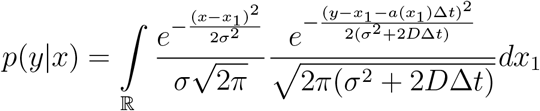. To obtain an explicit expression of this convolution, we use the change of variable *x*_1_ = *x* + *ση* where *σ* ≪ 1,

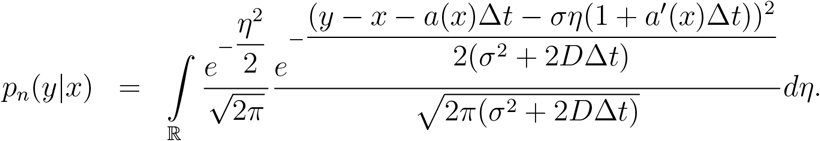

This integral can be regarded as the convolution of two gaussian functions over the real line, and

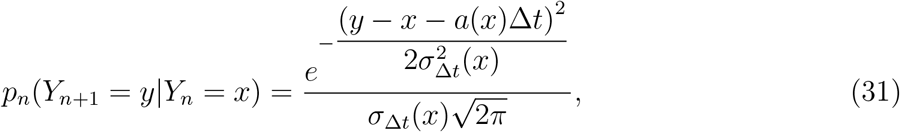

where

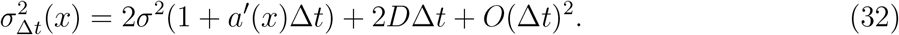

We conclude that the transition probability of the observed process *Y_n_* is Gaussian and 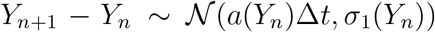. The observed motion is thus defined by the discrete scheme:

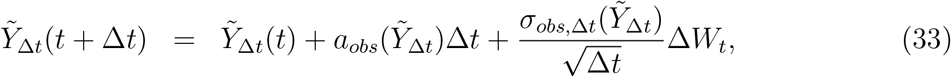

where Δ*W_t_* = *W*(*t* + Δ*t*) – *W*(*t*) and *W* is a Brownian motion of variance 1 and

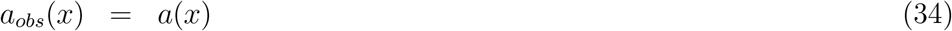

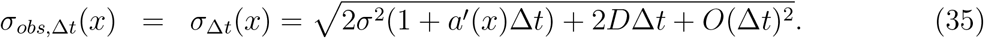

This approach allows defining the continuous process 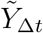 from the approximation at the scale Δ*t*, it is solution of the stochastic equation

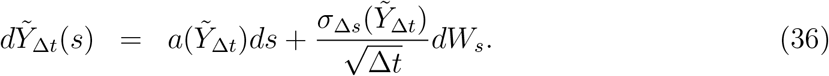

The drift of the observed process at a time resolution Δ*t* is the same (at first order in σ) as the physical one, while the diffusion tensor is changed and given by formula 32.

An estimator for the drift at a time resolution Δ of the observed process is constructed from

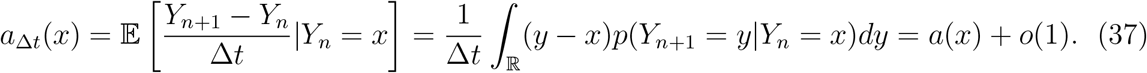

The result 27 is recovered: adding a Gaussian noise on the physical process does not alter the physical deterministic drift at first order in *σ*, while the diffusion coefficient is

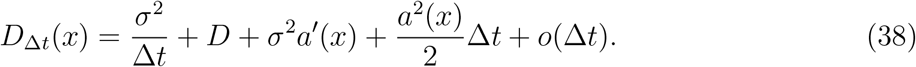

### 4.2 Generalization in dimension *m*

For a *m*-dimensional stochastic process

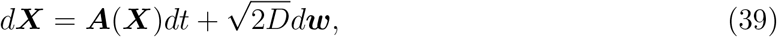

where ***A*** is a vector field and ***w*** the classical *m*–dimensional centered Brownian motion of variance 1. The diffusion tensor is assumed to be a constant *D*. The observed motion is

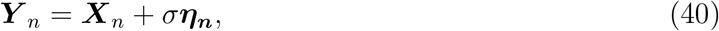

where ***η_n_*** are i.i.d *m*-dimensional standard gaussian variable and *σ* is a small parameter. Then, the estimator for the drift is [41]

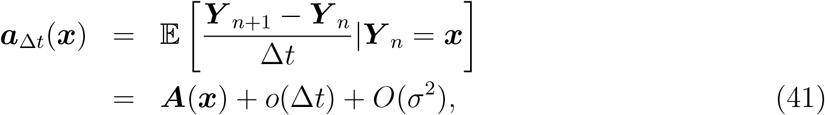

and the estimated diffusion coefficient relates to the physical parameter by

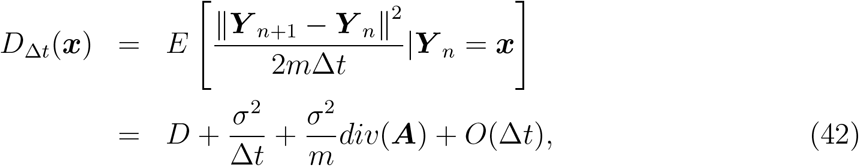

where *div*(***A***) is the divergence of the drift vector. These formulas show how spatial variations of the drift affect the measured diffusion tensor.

### 4.3 Other estimators

For a stochastic process containing a drift component, it is not possible to extract the physical diffusion coefficient directly by combining the first and the second moment estimators, which is in contrast with the pure diffusion case (see [6, 87]). We now present an estimator where the Gaussian instrumental noise can be eliminated. Using the difference Δ*Y_n_* = *Y*_*n*+1_ – *Y_n_*, we can rewrite

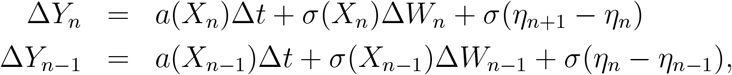

where Δ*W_n_* and Δ*W*_*n*−1_ are two independent increments of Brownian motion. The expectation is

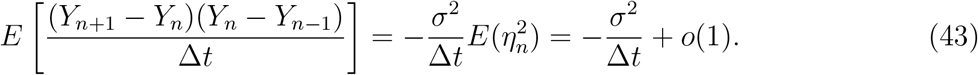

Using relation 38, we obtain that

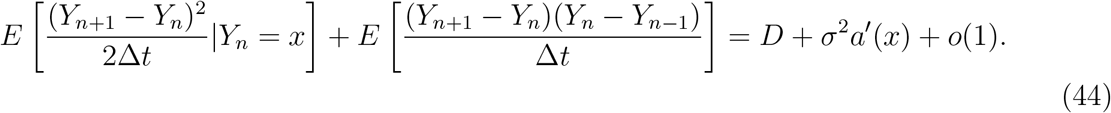

In this estimator, the instrumental noise is averaged out. There are no direct procedures to get rid of the derivative of the drift term, which can be extracted from the first order moment. However, computing a derivative from noisy data should be done carefully as it introduces singularities and irregularities.

### 4.4 Recovering the diffusion tensor in dimension 1

Using formula 38, we estimated the diffusion coefficient 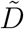 in Figs. 6A and 6B. The signal to noise ratio (SNR) is defined as 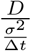. A high SNR can be due either to a large sampling rate Δ*t* or to a low positional noise. In our numerical application, we first vary the SNR by fixing the amplitude of the noise *σ* and by varying the increment Δ*t* (Figs. 6A and 6C) then we vary the parameters the other way around (Figs. 6A and 6D). We also estimated the diffusion coefficient for an OU process (Figs. 6B and 6D). These numerical estimations show that the estimator for the diffusion coefficient is biased for a high SNR for an OU process when the positional noise *σ* is fixed and the time step Δ*t* increases (Fig. 6C). This counterintuitive result is due to the approximation 23 for the physical motion 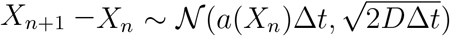, which is only applicable at small time steps Δ*t*. However this approximation is perfectly valid for a Brownian motion, as shown in Fig. 6A.

**Figure 6:**
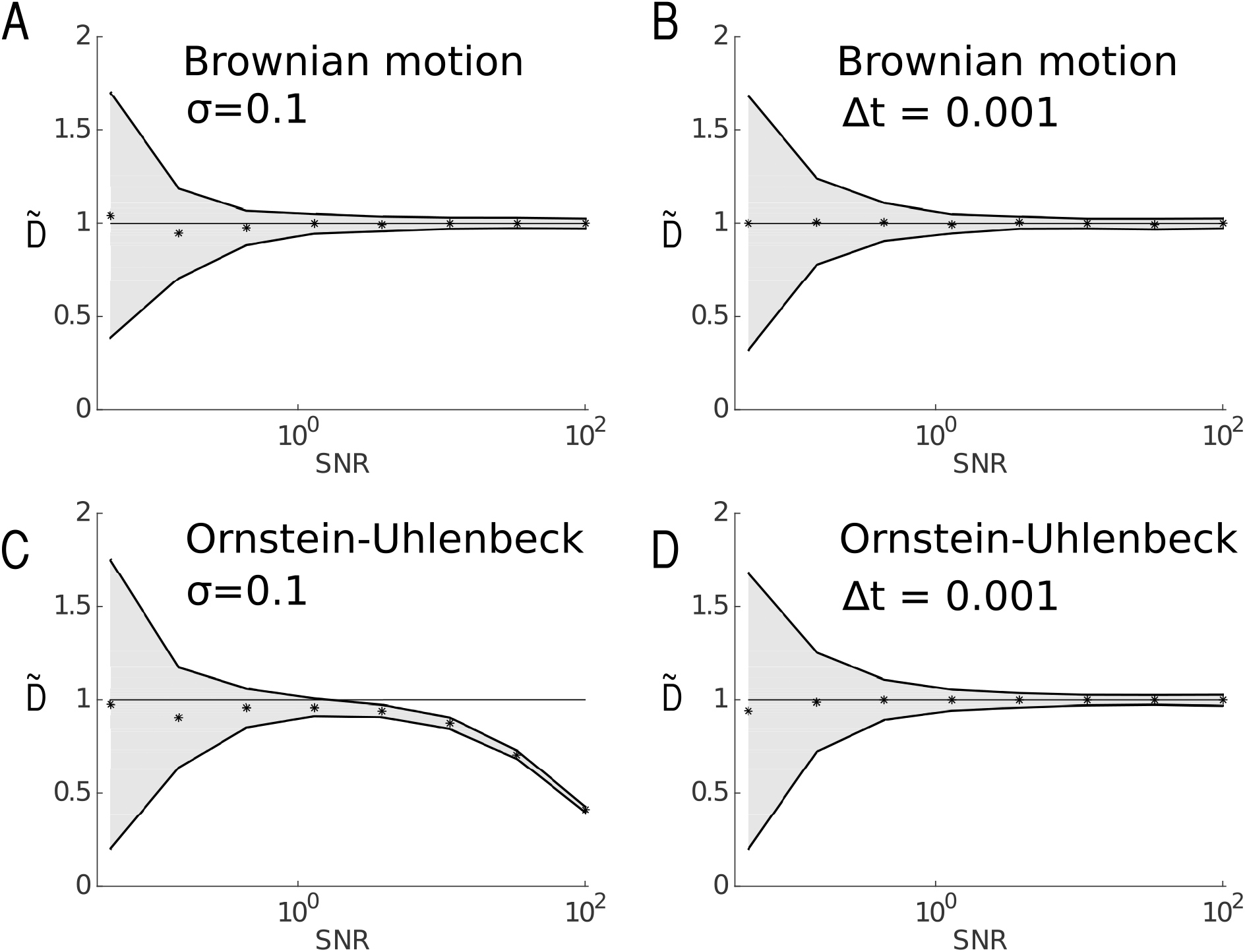
Diffusion coefficients are estimated for a Brownian motion (A and B) and Ornstein-Uhlenbeck process (C and D) and for various values of the signal-noise ratio (SNR) represented in a log-10 scale. Trajectories were simulated using Euler’s scheme, and sub-sampled so that the observed trajectories contain 10,000 points, and position noise *σ* was subsequently added to each point of the trajectories. The diffusion coefficient *D*_Δ*t*_ is estimated using formula 10. Black dots represent 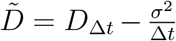 and the continuous lines bounding the grey area represent 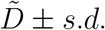. The diffusion coefficient is *D* = 1 and for the OU process the drift is *a*(*x*) = –2*x*. Variations of the SNR, defined as 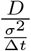, are obtained for fixed position noise (A,C) or fixed sampling time (B,D). In (A) and (C), the positional noise is fixed at *σ* = 0.1, while the sampling rate Δ*t* is varying. In (B) and (D), the sampling time is fixed to Δ*t* = 0.001 and the position noise *σ* is varying.

### 4.5 Empirical estimators associated to an Ornstein-Uhlenbeck process

For an Ornstein-Uhlenbeck (OU) process, 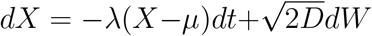, the pdf is 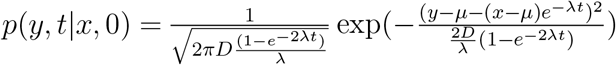. For the observed motion *Y_n_* given by eq. 30 the pdf is

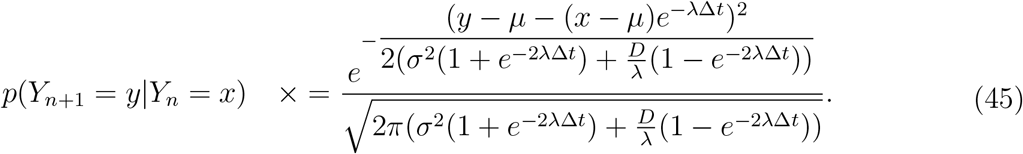

and the observed drift and diffusion coefficient at timescale Δ*t* are [41]

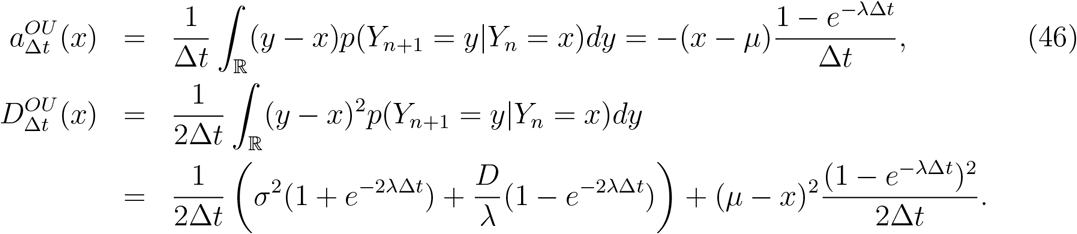

In Fig. 7, we estimate the local drift and diffusion coefficient for an OU-process and compare the local estimators for the drift 37 with relation 46 (Fig. 7B). For the diffusion tensor, we compare relations 38 and 47 (Fig. 7C). Using the first order approximation for short time step Δ*t*, we compare estimators 37 (resp. 38) that gives improved results compared to 46 (resp. 47), showing the improvement of the *O*(Δ*t*). Note that other classical statistical methods have been developed to recover drift and diffusion tensor [65, 1, 30] based on Bayesian inference or direct empirical sums.

**Figure 7:**
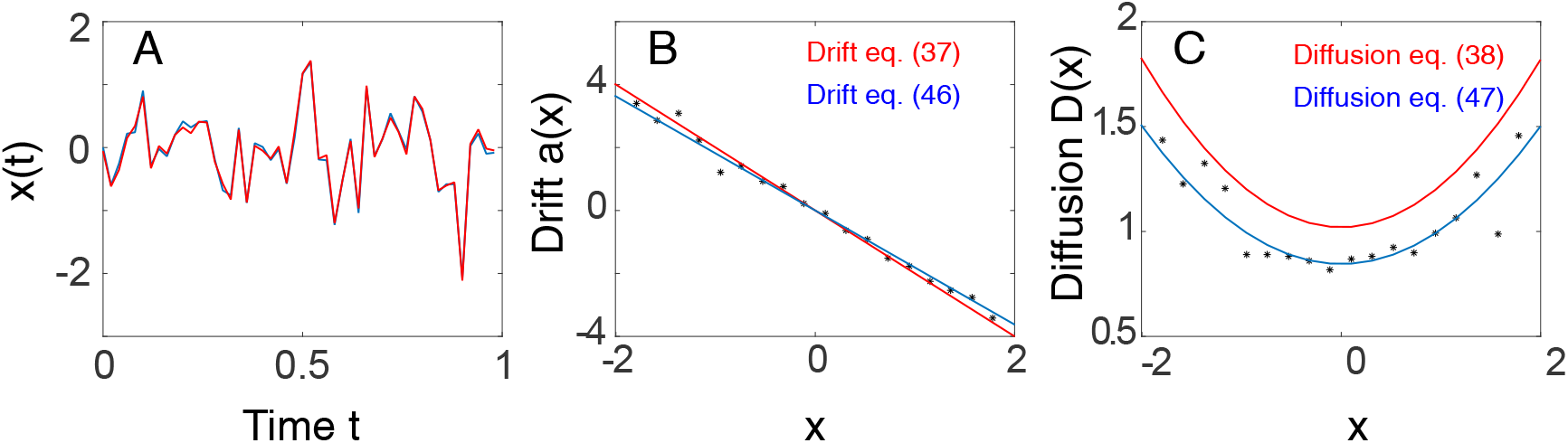
Recovering an Ornstein-Uhlenbeck process at second order in Δ*t*. (A) Trajectory of a one dimensional OU-process, generated using Euler’s scheme (blue curve) and the observed trajectory (red curve) is obtained by subsampling at Δ*t* = 0.1 and with an additional position noise of standard deviation *σ* = 0.05 (SNR=40). The other parameters are fixed to *D* =1, λ = 2, *μ* = 0. The observed trajectories contain 10,000 points. (B) Estimation of the local drift using eq. 9 (black dots), and comparison with the analytical formulas 37 (red) and 46 (blue). (C) Estimation of the local diffusion coefficient using eq. 10 (black dots) and comparison with the analytical formulas 38 (red) and 47 (blue), confirming the effect of the *O*(Δ*t*) term.

### 4.6 Maximum likelihood estimators for an Ornstein-Uhlenbeck process using the approximated and exact transition probabilities

We now describe briefly the maximum likelihood estimators to recover an OU process from the approximated transition probability. For a OU-process sampled at short time step Δ*t* (eq. 23) with the approximated scheme 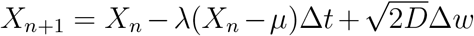, the transition probability of the observed motion is

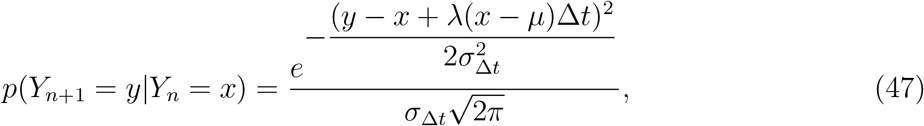

where the variance is

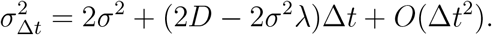

The log-likelihood is defined as

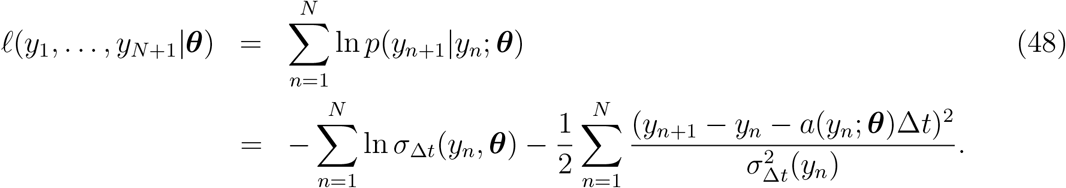

The likelihood function is applied here on a partially reconstructed OU-process, where the corrupted noise amplitude *σ* has already been accounted for in the variance term *σ*_Δ*t*_. Thus the discretization of the nonlinear SDEs for small time step Δ*t* does not rely on any addition assumption about the initial non-Markovian process of the observed measurements (generated from a discretize OU-process with an added Gaussian noise). The likelihood function 48 expresses the joint distribution of the observations as a product of transition densities, which is in fact valid for Markovian processes (i.e. a time series recorded without measurement error). But the observed measurement noise sequence is not a Markovian process. The exact likelihood would in principle requires using the entire series *p*(*y*_*n*+1_|*y_n_*; *y*_*n*−1_‥ 1;‥; *y*1; ***θ***). To overcome this difficulty, the effect of the noise is taken into account directly in the term *σ*_Δ*t*_. This procedure approximates a non-Markovian by a Markovian process. The variance term is recovered, however the error in the drift is not accounted for (see relation 27). In general, for an OU-process with constant measurement noise, the exact computation is tractable and detailed are given in ([30] p. 386). Yet, as we shall see the maximum likelihood estimator allows recovering the parameters of interest.

The parameters *θ*_1_,…, *θ_m_* and *D* are computed as maximizers of the likelihood function and thus by differentiating *ℓ* with respect to the variables *θ*_1_,…, *θ_m_, D*. The conditions 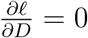 and 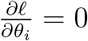 can be rewritten as

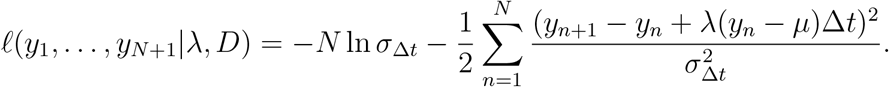

Conditions 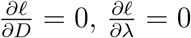 and 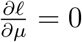 lead to

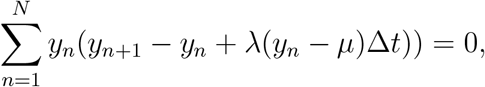

and thus the empirical estimator 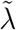 for the parameter *λ* is

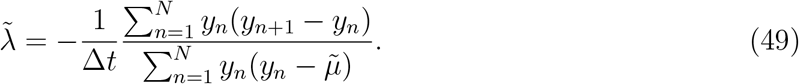

Similarly, using 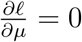, we obtain the condition

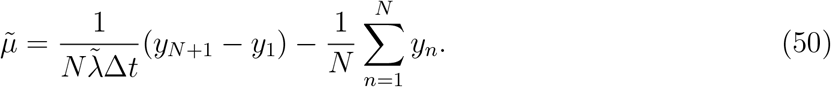

By combining 49 and 50 we obtain

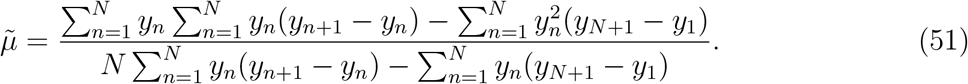

Finally, the empirical estimator for the diffusion coefficient is

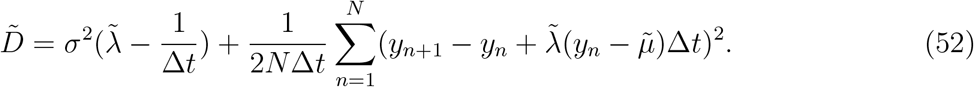

Estimators can also be built using the exact transition probability of the OU process

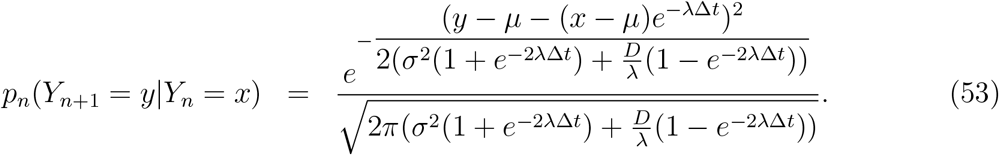

and are given by [41]

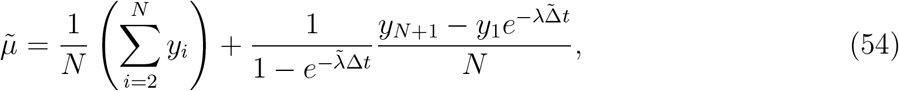

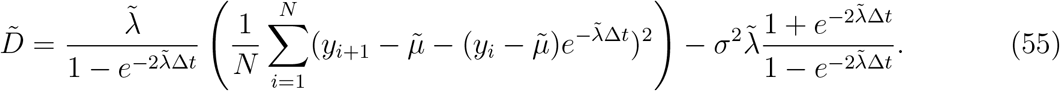

The two maximum-likelihood estimators for the OU process are compared using numerical simulations, where the time-step is Δ*t* = 0.1 and the parameters *λ, μ* and *D* are estimated for *n* = 500 observations. The parameters are estimated for various values of the observation noise *σ*. The results are summarized in Fig. 8: the parametric estimators based on the exact transition probability of the OU process, gives better estimates than for the one using the approximated pdf. To estimate the likelihood function, other approaches are based on Kalman filtering [30].

**Figure 8:**
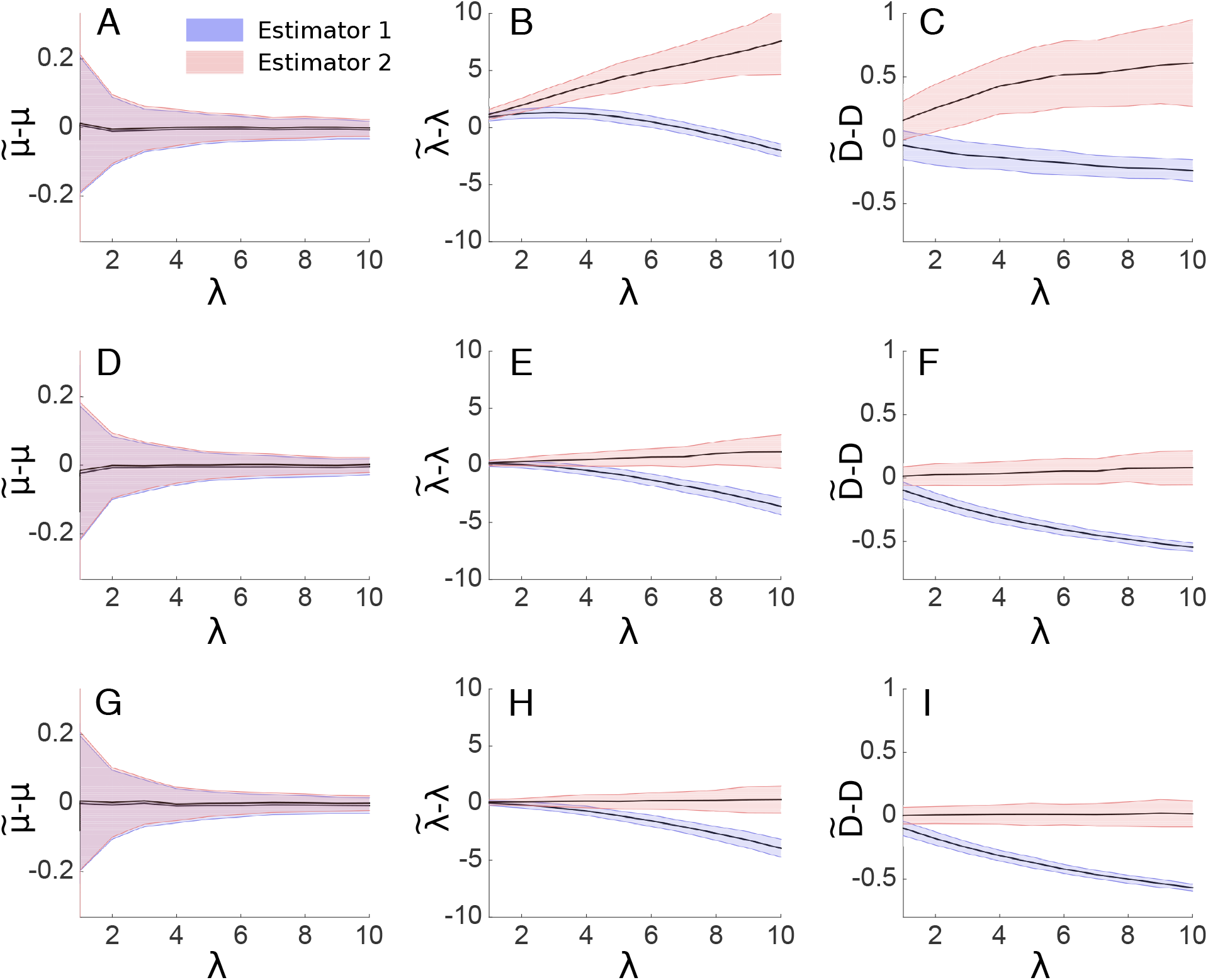
The maximum-likelihood estimators presented in 4.6 (blue: approximated pdf, red: exact pdf) for an OU process with *D* = 1, *μ* = 1, and various values of λ are compared. From left to right, the plotted estimations are 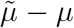 (A,D,G), 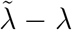 (B,E,H), and 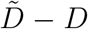 (C,F,I). The observation time-step is Δ*t* = 0.1 and from top to bottom, 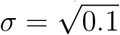 ((A,B,C), SNR=1), 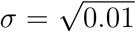 ((D,E,F), SNR=10), 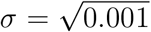 ((G,H,I), SNR=100). The black line is the mean of the estimation for trajectories of 500 points, and the colored part indicates *±* the standard deviation (adapted from [41])

### 4.7 Parameter estimation for motion with varying dynamics

For non-stationary processes, a time correlation between points of SPT trajectories renders the analysis quite difficult: indeed when particles can alternate between free and confined motion, or experience rapid changes in the interacting forces [12, 88], the process is not stationary. Similarly, potential wells can appear or disappear [40] at a time scale of minutes. To extract information from non stationary SPT trajectories, the Hierarchical Dirichlet Process Switching Linear Dynamical System (HDS-SLDS) method [20, 21] detects abrupt changes in the forces governing a single trajectory [11] (see also [9, 12]). The method extends hidden Markov models (HMM) [83], in which the different states are associated with a linear dynamical process. The underlying discrete physical model of the motion is

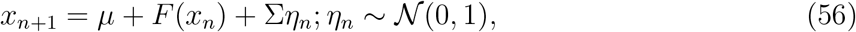

where *x_n_* is the position at the discrete time step *t_n_, μ* is a constant vector, *F* is a space-dependent force matrix, and *η_n_* a random variable to account for fluctuations in the particle motion. The observed position is the sum of the physical position plus a normal random variable with variance *σ* (localization noise)

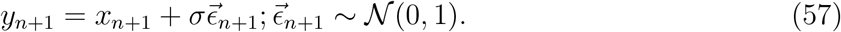

When the motion alternates between different states, characterized by the parameters *θ* = (*μ, F*, Σ, *σ*), the procedure consists in decomposing a trajectory in different segments using a non-parametric Bayesian technique. The motion parameters in each sub-trajectory is estimated using a maximum likelihood estimation (MLE) method. The transition between each state is detected as large jumps in consecutive points of the observed trajectory.

In the framework of the HMM, transitions between the different states are modeled as Markovian probabilities. Finally, to classify noise terms generated by thermal motion, external forces or the localization noise, a Kalman filter can be used to estimate the physical position of the particle. This method was used to detect various regimes in chromatin motion in yeast cells and to estimate parameters in each cell stage [11].

### 4.8 Comparison with classical single particle trajectories analysis

The ensemble of estimators described above are based on the exact expression computed from the statistical moments and pdf of the stochastic processes. We now recall the classical mean-square displacement (MSD) estimators. The diffusion coefficient is computed from single particle trajectories using the MSD, which consists of averaging the square displacement over single trajectories: in one dimension, for a sequence of points (*x*_1_,…, *x*_*N*+1_) sampled at constant time intervals Δt, the MSD is given by [64]

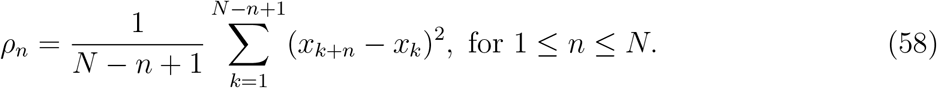

For a Brownian particle with a diffusion coefficient *D*, the MSD is given by

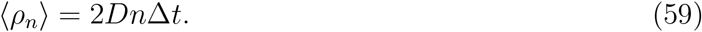

This relation between the MSD and the diffusion coefficient is however not a good estimator of the diffusion coefficient for a general stochastic process that contains a drift or a localization noise, but also for a Brownian motion [6, 87, 88, 16]. For example, the accumulation of the noise by adding points along the trajectory affects the extracted value of *D*. The typical MSD curve differs from a line because at large time lags, the number of points used in the averaging is very low and the squared displacements are not independent from each other, resulting in cumulating errors [64]. Moreover, this procedure does not preserve the spatial heterogeneity of regions explored by the random particle.

The finite exposure time (see relation 19) affects the estimate of the diffusion coefficient for particles undergoing hop diffusion or confined motion [67]. The measured diffusion coefficient *D* decreases as the exposure time increases. The motion blur has even more effect on fast moving particles, as the particle observed position is the result of integration over longer areas. Including motion blur in the MSD leads to the following estimate for MSD [88]

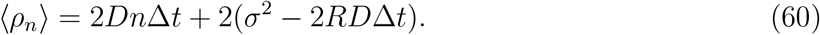

In the case of a motion blur and position noise, optimal estimates of the diffusion coefficient uses an optimal number of points in the MSD formula [59]. Localization error and the finite camera exposure affect the MSD curve and thus perturb the estimation of the diffusion coefficient [72, 61, 28]. In the absence of localization error, the optimal estimate of the diffusion coefficient uses the first two points of the MSD [74, 59]. To conclude, fitting the MSD curve is difficult because the sequence *ρ_n_* are highly correlated. The estimation of the diffusion coefficient becomes imprecise as the number of points included in the fit increases [59]. The MSD remains commonly used [64] to estimate the diffusion coefficient despite localization and dynamical noise that alter significantly the MSD slope at short timescales, which might be erroneously attributed to subdiffusion processes [6].

## 5 Statistical analysis of a trajectories from a single locus located on a polymer

Long SPT trajectories can be used for extracting biophysical parameters and mechanical forces. Indeed, a dye molecule located on a DNA is often used as a probe to explore the local cellular nucleus environment. We present here several estimators used to reconstruct external and internal forces applied on the DNA molecule and chromatin from the statistics of long locus trajectories. In the stochastic model, the locus motion is driven by local diffusion plus forces between monomers of a model polymer [45, 89, 86]. Monomer motions are usually highly correlated leading to anomalous diffusion behavior [57, 42].

To distinguish external forces applied on a single monomer from intrinsic forces acting on monomers, the first step is to use a polymer model such as Rouse. A stochastic particle is modeled by the Smoluchowski’s limit of the Langevin equation: the velocity **v** is proportional to a force **f** plus an additional white noise, summarized as

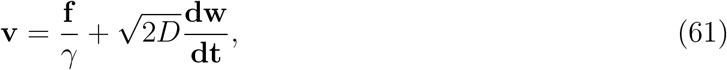

where *γ* is the friction coefficient, *D* the diffusion coefficient and *w* is the normalized Wiener process. By averaging over the ensemble of velocity realizations, the first moment or the force field is recovered [37]. However, for a polymer chain, there are internal forces between monomers. In recent statistical methods, it was possible to separate from measured single (monomer) trajectories, internal polymer forces from the external ones acting on a single localized monomer.

For an external force which is the gradient of the potential *U*_ext_(***R***) and applied to a Rouse polymer, the physical interactions are described by the energy term

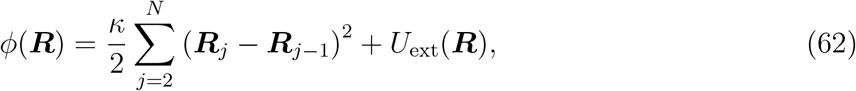

where ***R*** = (***R***_1_, ***R***_2_,…, ***R***_*N*_) is the ensemble of monomer positions, connected by a spring of strength *κ* = *dk_B_T*/*b*^2^. *b* is the standard-deviation of the distance between adjacent monomers [17] and *d* the dimensionality (dim 2 or 3). In the Smoluchowski’s limit of the Langevin equation [77], the dynamics of monomer ***R***_*n*_ is described by

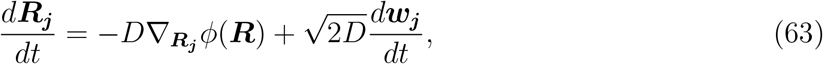

for *j* = 1,…, *N* and each ***W**_j_* is an independent *d*-dimensional white noise with mean zero and variance 1, *D* is the monomer diffusion coefficient. We will describe specifically the field of force ∇_***R**_j_*_ *U*_ext_(***R***) in the next subsection.

A reference monomer ***R**_c_* is chosen representing a tagged locus and the first statistical moment of the monomer ***R**_c_* (averaged over all realizations) is recovered from data from an inversion formula, generalizing eq. 8:

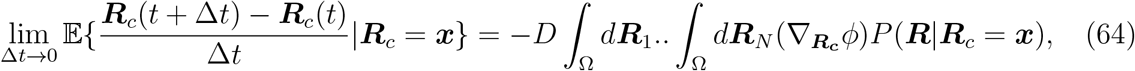

where 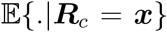 denotes averaging under the condition that the tagged monomer is at position ***R**_c_* = ***x***. Formula 64 is generic and does not depend on the external forces acting on the polymer.

When a polymer chain evolves in a domain Ω, with reflecting condition on the boundary *∂*Ω, the conditional probability *P*(***R***|***R***_*c*_ = ***x***) is computed from the equilibrium probability distribution function *P*(***R***_1_, ***R***_2_,…, ***R***_*N*_), which satisfies the Fokker-Planck equation (FPE) in the phase-space 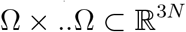,

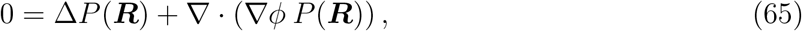

with boundary condition

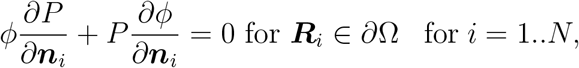

where ***n**_i_* is the normal vector to the boundary *∂*Ω at position ***R**_i_*. A permanent force located at position ***μ*** can be approximated by an harmonic well. A force is applied to monomer *n* located at position ***R**_n_* and is computed from the gradient of the harmonic potential

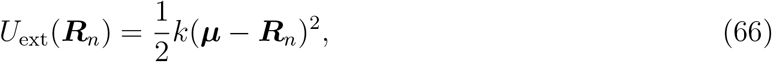

where *k* is the force constant. Any monomer *n* is affected by the applied force, but differently from the tagged monomer *c* and we shall assume that *n* < *c*. The potential well affects the dynamics of the entire polymer and specifically the observed locus *c*. For the mean velocity of monomer *c*, we get the following relation

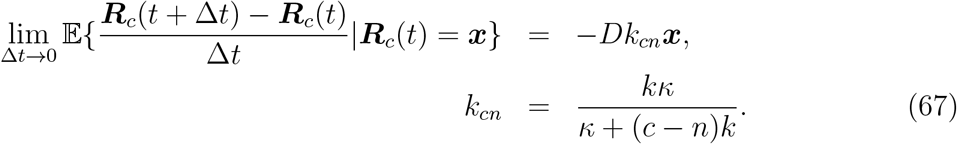

Expression 67 links the average velocity over empirical trajectories of the observed monomer *c* to a permanent force applied on monomer *n*. The coefficient *k_cn_* depends on the harmonic well strength *k*, the inter-monomer spring constant *κ* and is inversely proportional to the distance |*n* – *c*| between monomers *n* and *c* along the chain.

To extract the empirical effective spring coefficient *k_c_* from a trajectory given by ***R**_c_*(*h*Δ*t*) (*h* = 1… *N_p_*), where *N_p_* is the number of points, the differential quotient in eq. 67 is first computed. This step allows extracting the mean position of the locus. Once the steady state is reached, the time average of the locus position is computed from

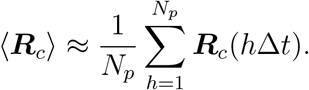

When the polymer interacts with a single interacting potential, the average position 〈***R_c_***〉 estimates the location where the force is applied. An upper bound for the number of points *N_p_* is of order (*τ_r_*/Δ*t*), where 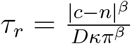 is the relaxation time for a portion of the chain of a *β*-polymer between monomers *c* and *n* [2].

When the diffusion coefficient *D* is known (which could be extracted using the second moment estimator 10) and the inter-monomer spring constant *κ* are known, it is possible to estimate the force from the constant *k_cn_* in eq. 67, using the linearity of the force with respect to the position of the locus (see eq. 66). The step size (***R***(Δ*t*(*h* + 1)) – ***R***(Δ*th*)) is proportional to the locus position (***R***(Δ*t*(*h* + 1)) – 〈***R**_c_*〉). In the isotropic case, the apparent force constant *k_cn_* acting on monomer *c* is computed from the trajectories of ***R**_c_*(*t*) by the estimator

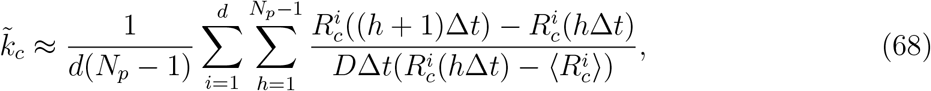

(*d* is the dimension and *N_p_* is the number of points). This approach has been validated using stochastic simulations and live cell imaging data [3]. For a strong anchoring (*k* ≫ *κ*), we can approximate *k_c_* ≈ *κ*|*c* – *n*|^−1^. In that case, the empirical effective spring constant can be used to estimate the distance to the interacting monomer.

For a long enough sampled trajectory and a force derived from a stationary potential well, the effective spring coefficient can be recovered directly from the empirical estimator 68. However, with a small sampling step Δ*t*, refined chromatin behavior can be recovered accurately as well as the diffusion coefficient. However, the length of a trajectory is often limited due to photobleaching [66]. In principle, the length of a trajectory may be shorter than the time scale to equilibrium. In that case, formula 68 can still be applied to recover the parameter *k_c_*.

## 6 Reconstruction of the surfaces of cell from the statistics of observed projected random trajectories [39]

A large samples of short trajectories of diffusing single particles acquired on the surface of a cell are projected onto the plane of a confocal microscope. How these projected data can be used to reconstruct the shape of the membrane surface and physical properties of the receptor motion? A general method for the reconstruction of a two-dimensional surface from the statistics of planar projections of many independent trajectories of a diffusion process on the surface introduced in [39] starts with the finding the original stochastic dynamics from the projections. The reconstruction of the geometrical properties and drift field, which represents physical interactions, falls into a class of inverse problems that have been investigated in the context of extraction of shape from shading [90].

However, due to the irregularity of stochastic trajectories, the reconstruction method requires in practice a large set of data points. The reconstruction scheme is illustrated in Fig. 9. The driftless case can be formulated as follows. The planar projection ***Y*** of a diffusion process ***X***, defined on a smooth manifold Σ by the Itô system

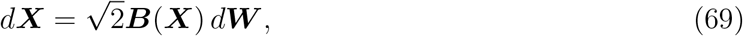

where ***W*** is Brownian motion on Σ and ***B***(***X***) is the noise matrix, satisfies the equation

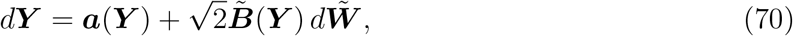

where ***a***(***Y***) and ***B***(***Y***) can be expressed in terms of the derivatives of Σ on the projection plane (e.g., a fixed tangent plane 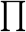). Because Σ is unknown, ***a***(***Y***) and ***B***(***Y***) are the effective drift field and diffusion matrix observed in the plane. Thus the first task is to obtain an analytic expression for these functions. Once explicit expressions for the effective drift field and diffusion tensor in terms of the local surface properties such as the mean curvature are found, the next task is the reconstruction of the surface, which requires solving nonlinear partial differential equations. When the dynamics on the surface contains a drift field, the drift field of the projected dynamics is split into a part that is due to the surface curvature and another due to the original drift.

**Figure 9:**
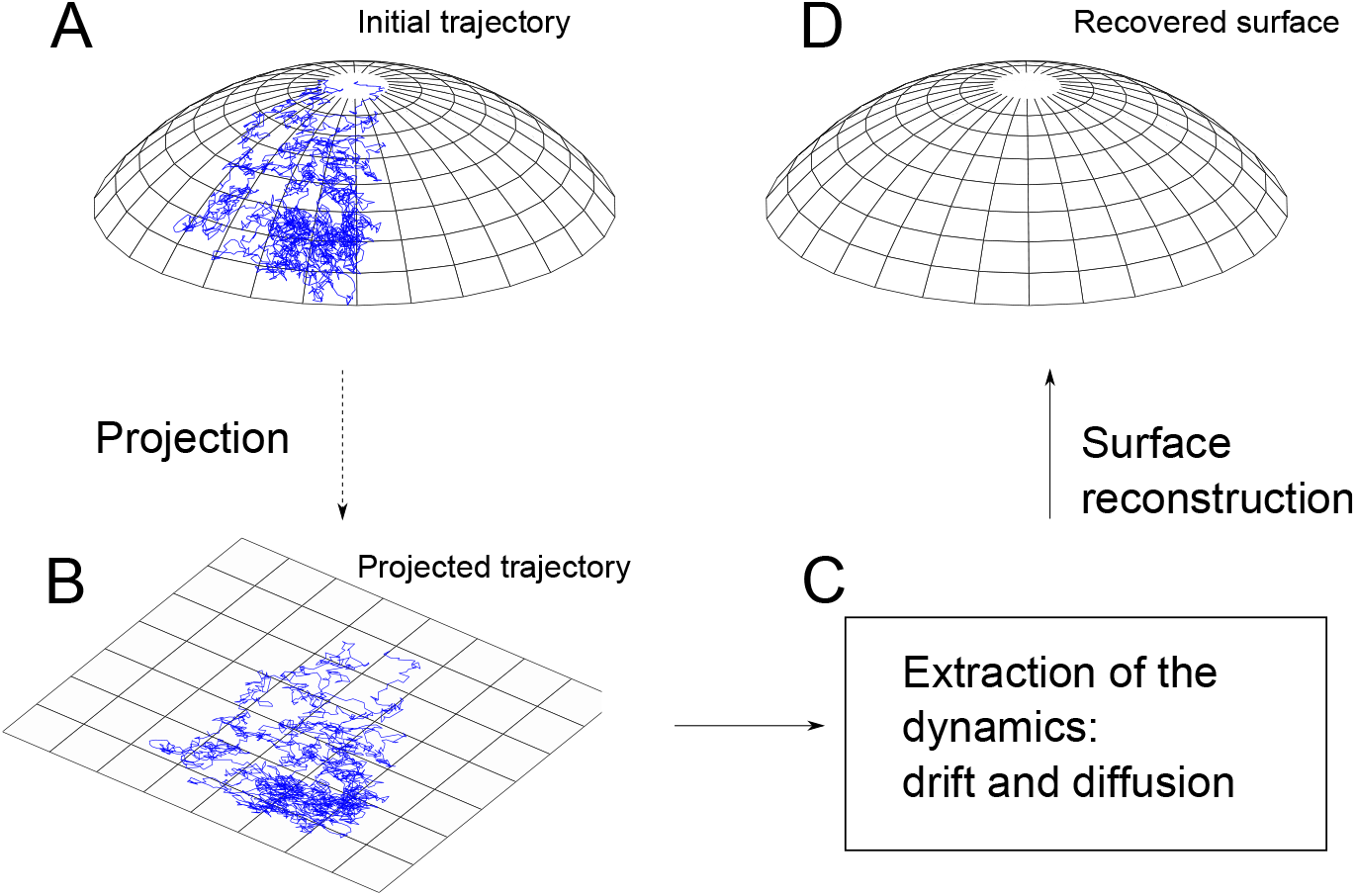
Reconstruction of a surface from the dynamics of a projected trajectory. (A) A trajectory of a diffusion process on the unknown surface Σ. (B) A planar projection of the trajectory. (C) Statistical estimates of the drift and the diffusion coefficients of the projected stochastic dynamics. (D) Reconstruction of Σ from the estimated coefficients.

We present here the main step of the reconstruction for a Brownian motion ***X***(*t*) on a surface Σ and its planar projection. We assume that Σ has the explicit representation *z* = *f*(*x,y*), where *f*(*x,y*) is a sufficiently smooth function defined in a planar domain 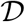 in the (*x, y*) plane. We can assume that the (*x, y*) plane is tangent to Σ at the origin **0**, that is, **0** £ Σ and the normal to Σ at **0** is the *z*-axis. We fix an orthonormal frame (***i, j, k***), where ***k*** is the unit vector in the direction of the *z*-axis. Thus

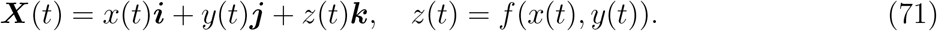

For a driftless diffusion ***X***(*t*) with a symmetric diffusion tensor *σ^i,j^*(***X***) on Σ with a tangent plane 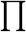 at ***X***(*t*) ∈ Σ, we define the orthonormal frame

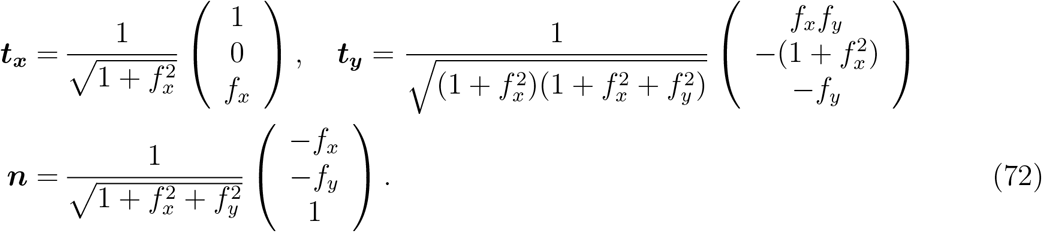

The orthogonal projection 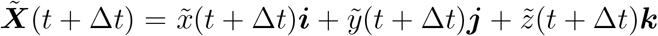 on 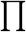 of the Brownian motion ***X***(*t* + Δ*t*) ∈ Σ is given in terms of 72 by

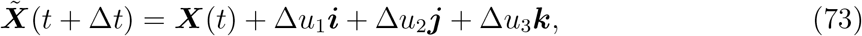

where

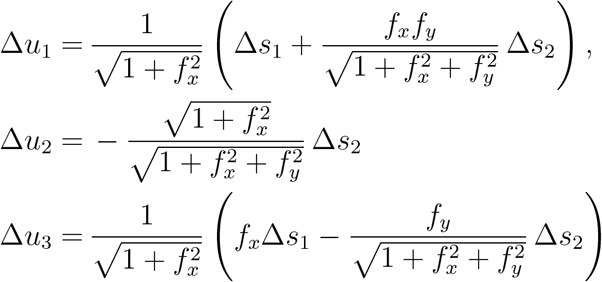

and

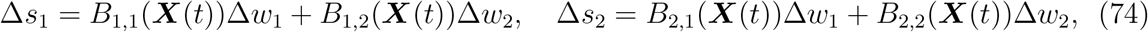

with *w*_1_(*t*), *w*_2_(*t*) independent standard Brownian motions in 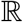. The diffusion tensor ***σ***(***X***) is given in terms of the matrix ***B***(***X***) as

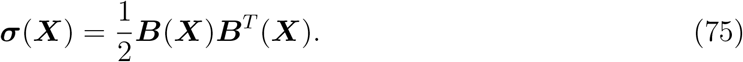

To regain ***X***(*t* + Δ*t*), the point 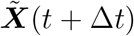 is projected orthogonally back from 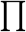 to Σ to give

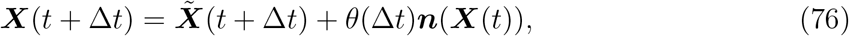

where the parameter 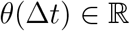 is the solution of the equation

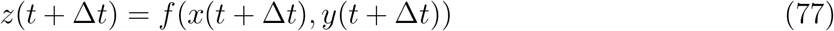

with

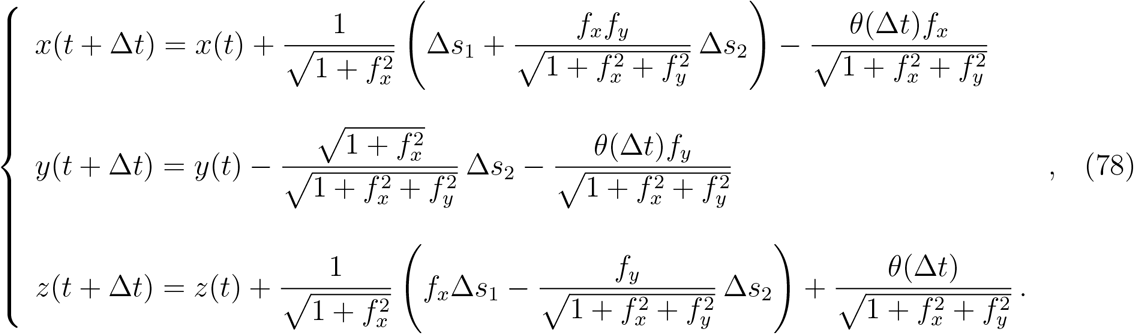

where *θ*(Δ*t*) is such that [39]

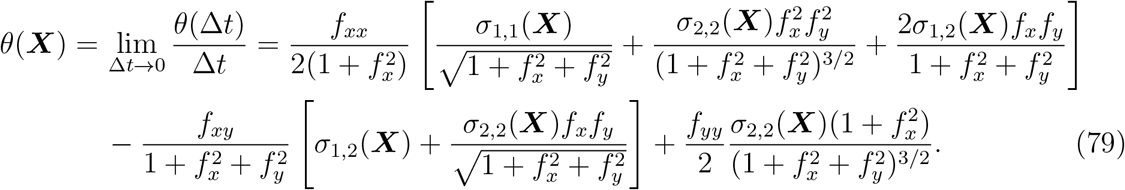

In the isotropic case ***B***(***X***) = ***I***, we obtain

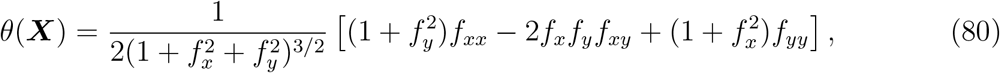

which is the mean curvature at point ***X***. Setting

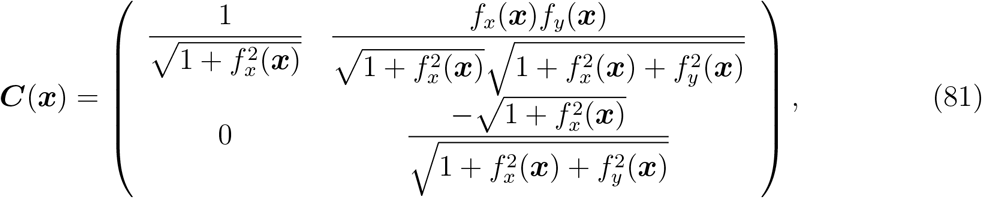

we define

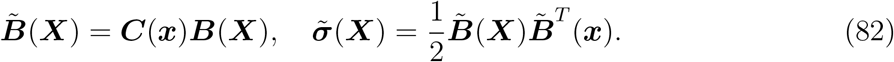

Thus the system of stochastic equations in 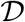 that describes the driftless diffusion *z*(*t*) = *f*(*x*(*t*), *y*(*t*)) on Σ, where the projected process on the (*x,y*) plane, ***x***(*t*), is

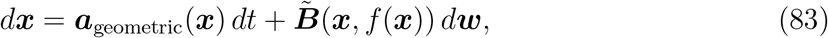

where

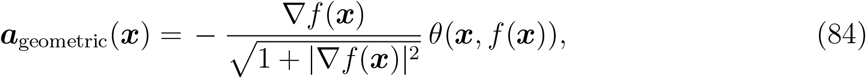

and ***w***(*t*) = (*w*_1_(*t*), *w*_2_(*t*))^*T*^. If 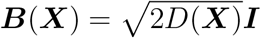, then

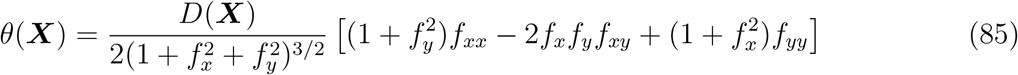

and 83 becomes

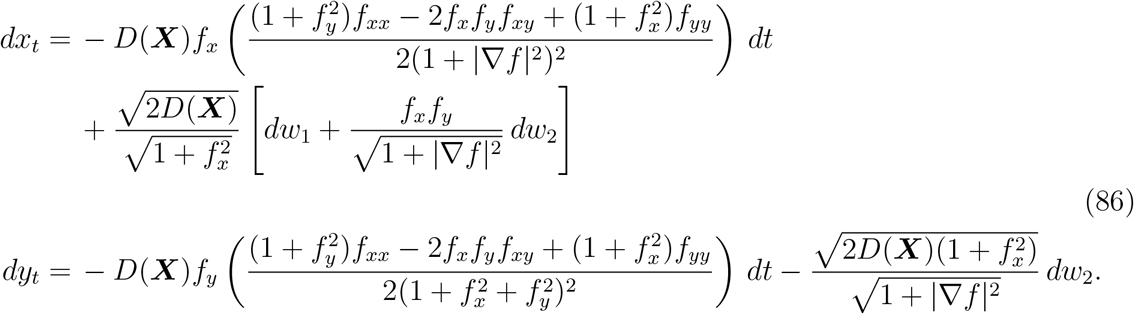

The planar projection of the Brownian motion (or driftless diffusion) has a geometric drift vector and an effective diffusion tensor that are observed. The surface Σ and the diffusion tensor can be recovered from the observed projections.

When the diffusion process ***X***(*t*) on Σ has a drift vector ***β***(***X***) = *β_x_**i*** + *β_y_**j*** + *β_z_**k***, such that *β_z_*(***X***) = *β_x_*(***X***)*f_x_*(***x***) + *β_y_*(***X***)*f_y_*(***x***), the stochastic equations of the projected diffusion contain an additional drift vector

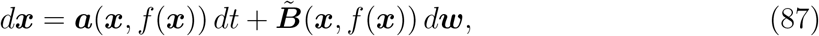

where

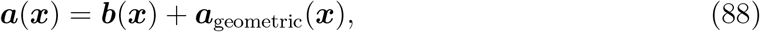

with ***b***(***x***) = (*β_x_*(***X***); *β_y_*(***X***)).

### 6.1 Reconstruction procedure

For an isotropic constant diffusion, the projection of Brownian motion from Σ to the (*x, y*) plane and recover the function *z* = *f*(*x, y*) and the diffusion coefficient *D* from the effective diffusion tensor

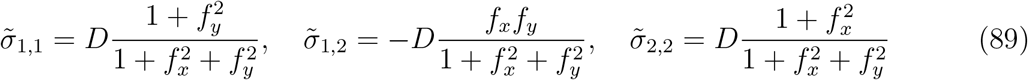

given in 82. The system 89 of three algebraic equations for the unknowns *D, f_x_*, and *f_y_* can be solved explicitly and then Σ is recovered by integrating the gradient ∇*f*(*x, y*) and reduces to solve the single equation new Eikonal equation

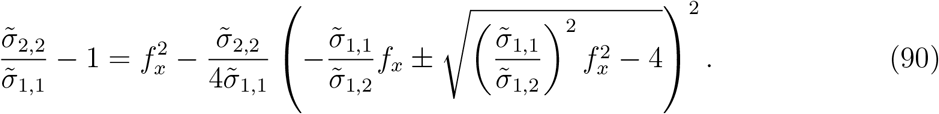

Before recovering the drift, because

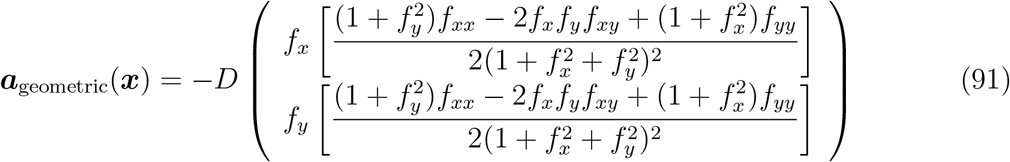

equation 91 implies that ***a***_geometric_(***x***) ║ ∇*f*(***x***) so the level curves of *f*(*x, y*) can be determined by integrating the geometric drift at any point in the planar projection of Σ. To recover isotropic Brownian motion with constant diffusion coefficient on Σ from its planar projection, we determine the components ***b***(***x***) = (*b_x_*(***x***), *b_y_*(***x***))^*T*^ of the drift by solving the system 88, which can be written as

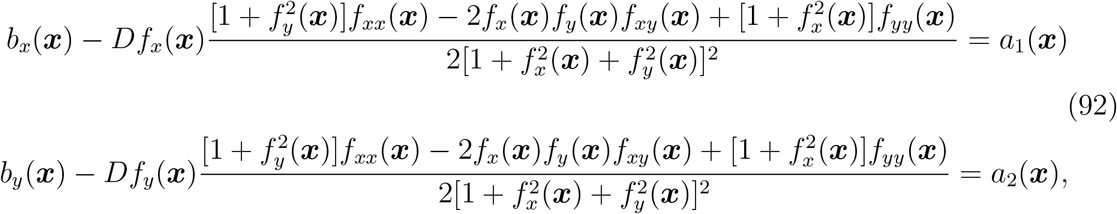

where now the functions *a*_1_(***x***) and *a*_2_(***x***) are the observed drift components of the projected diffusion. However, numerical reconstructions are not easy and require many points.

## 7 Conclusion and open questions

This manuscript presented a statistical approach to single particle trajectories, where estimators are constructed to extract first and second order moment properties of the underlying physical processes while removing various sources of noise. The large amount of SPT trajectories allows recovering drift and diffusion tensor at each spatial point at a given resolution. In contrast, computing the MSD along trajectories leads to accumulation of errors due to localization and instrumental noise and also correlates points in the MSD estimation.

Several improvements are expected in the future. For example, diffusion in confined domains has been widely studied in corrugated channels, in domains with obstacles, but the effect of confinement on the observed diffusion coefficient remains difficult to address in particular at the boundary with obstacles. This question is related to the finite time step which is often too large to resolve the molecular size and thus the physical diffusion. Extracting the radius of confinement [49] is very tempting, but in practice, confinement can also be due to an interacting potential. It is a key step to clarify the contribution of both effects and thus the first and second moments should be accurately computed and interpreted. Stochastic simulations and analysis reveal that a random walk has an apparent drift near a wall, which is proportional to 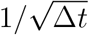, where Δ*t* is the sampling rate. This relation provides a criteria to differentiate hard-wall confinement from a deterministic drift, for which the estimator is independent of the sampling rate [41]. This criteria could be implemented in confined domains of size larger than the layer 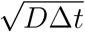. To conclude, a multiscale analysis with various Δ*t* should be used to resolve the presence of a boundary of a subdomain versus the confinement due to a potential well.

In general, how to integrate positional noise and motion blur on the estimate of diffusivity parameters in a confined domain is still an open question. Several studies [37, 55] have now confirmed that regions where trajectories are very confined are due to potential wells. However, extracting the exact properties of the wells remains difficult due to estimation errors of the boundary and the energy constant [63]. Another difficulty in interpreting the SPT trajectories in cell biology is due to the small size of the confinement regions in the range of 100 nm, which is of the order of the localization noise. Resolving hard-wall reflection from interaction potential would be a challenge for statistical methods, but is it really a biophysical issue? Probably hard-walls are a simplification of three dimensional scaffolding proteins at the interface between two- and three-dimensions, that should be addressed using the geometry of the molecules. Measuring and observing particle interactions [10] has recently been applied for G-protein coupled receptors forming dimers or oligomers [43]. But investigating more complicated motions and extracting parameters beside diffusion or interactions (modeled by an Ornstein-Uhlenbeck process) is usually more involved especially for processes driven by molecular aggregation-dissociation. In that case, there is no long-range interaction and thus no potential wells are expected. A statistical analysis is clearly missing.

Most estimators rely on the stationary assumption. Even when a particle motion alternates between free and confined motion [88], it is always possible to decompose the trajectories into sub-time windows, where the motion is clearly identified. This approach of time segmentation was applied to identify transient potential wells that appear and disappear [40], revealing that these structures are generated by dynamical processes, yet to be identified. Indeed, the analysis was performed for sufficiently small windows of time (of the order of seconds) and during that time, potential wells are stationary, but not at a time scale of minutes, where they can evolve [37]. It is always interesting to detect the different time scales involved in subcellular processes. For example, detecting events such as clustering of proteins during the formation of vesicle (clathrin-coated pits [51]) should certainly benefit from a statistical analysis to identify the exact mechanism of clustering.

The method for recovering a stochastic dynamics and the surface using the projection of short stochastic trajectories on a plane is not converging very quickly, because stochastic trajectories are not smooth enough and it is thus difficult to obtain an accurate reconstruction of the surface. A large number of points are necessary to recover surfaces. In dimension two, the sampling number of points is *N*^2^ (compared to *N* in dimension one). Using simulations of 9,000 trajectories, each composed of 1,000 points on a sphere and projected on the (*x, y*) plane, the effective diffusion tensor and geometric drift can be evaluated by formulas 9, 10 (Fig. 10A-C). The surface is reconstructed by integrating the gradient, as described in section 6.1 (Fig. 10F). Then the drift vector ***b***(***x***) = (*b_x_*(***x***), *b_y_*(***x***))^*T*^ of the projected diffusion is found by subtracting the geometric drift from the measured drift ***a***(***x***) = (*a*_1_(***x***), *a*_2_(***x***))^*T*^ (see (92)). This procedure is however not very practical.

**Figure 10:**
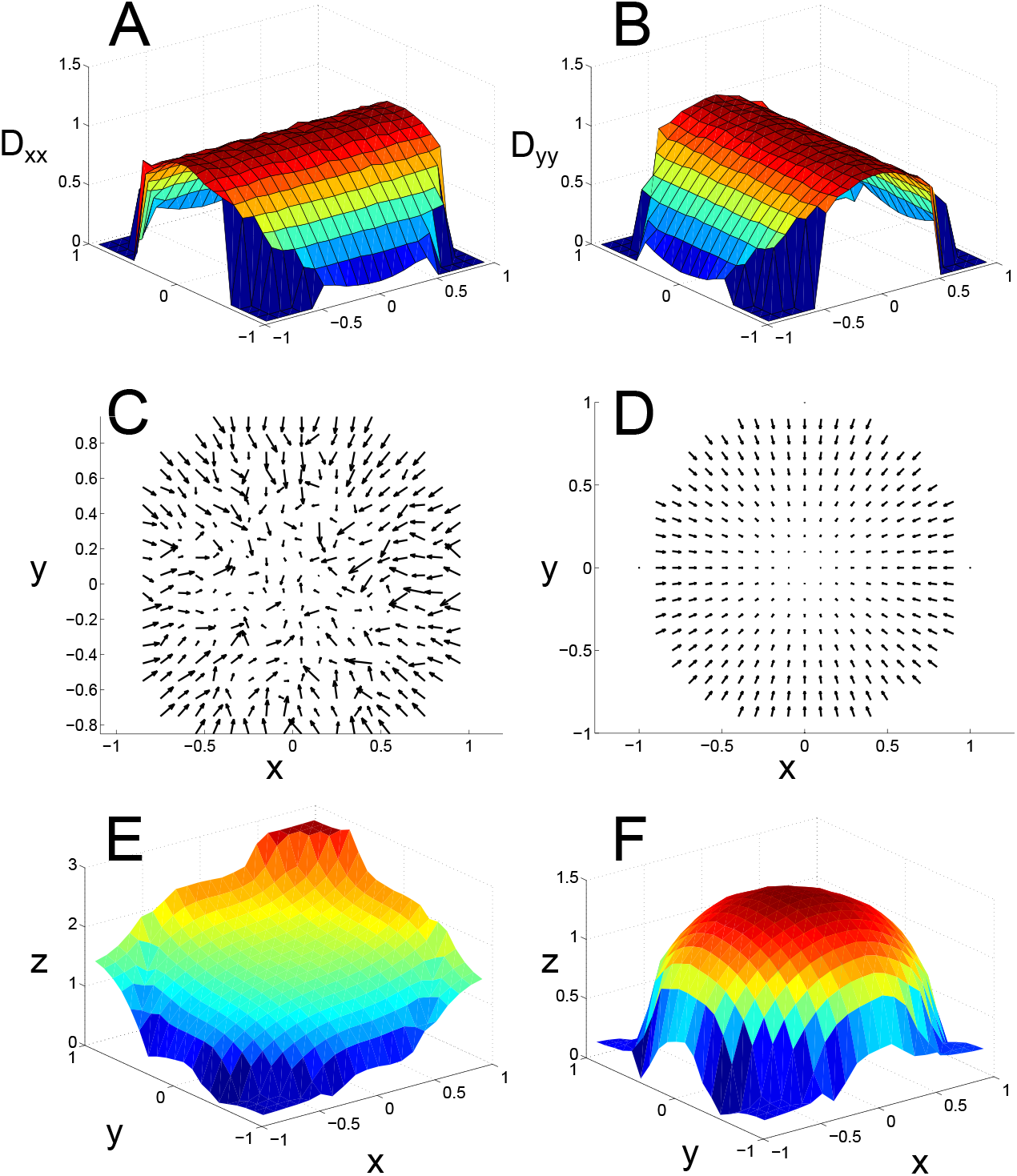
Projection of spherical Brownian motion. (**A**) Diffusion coefficient *D_xx_* reconstructed using equation 10. (**B**) Diffusion coefficient *D_yy_*. (**C**) The drift field ***a***(***x**_k_*) = (*a_x_*(***x**_k_*), *a_y_*(***x**_k_*))^*T*^, reconstructed by formula 9. (**D**) The theoretical drift field of the projected Brownian motion on a sphere, obtained from equation 86. (**E**) Reconstruction of the sphere assuming *f_x_* ≥ 0 and *f_y_* ≥ 0. (**F**) Reconstruction of the sphere. The parameter values are *D* = 1, *dt* = 0.00005, number of trajectories *N_t_* = 9, 000, number of points per trajectory *N_s_* = 1, 000, Δ*X* = 0.1.

Finally, three dimensional SPT trajectories will be soon available [60, 35], and an approximation theory should probably be used to reconstruct the dynamics of the underlying process and the membrane surface or the three dimensional cellular domain, where trajectories evolve. Statistical analysis should be used to improve the spatial resolution based on redundant trajectories, especially when the resolution in the z-direction is much less than in x and y. Probably, the sampling time Δ*t* will not change. Multiscale spatio-temporal data should further improve the reconstruction of cell surfaces.

